# Proteomic Architecture of Human Coronary and Aortic Atherosclerosis

**DOI:** 10.1101/157248

**Authors:** M Herrington David, Mao Chunhong, Parker Sarah, Fu ZongminG, Yu Guoqiang, Chen Lulu, Venkatraman Vidya, Fu Yi, Wang Yizhi, Howard Tim, Goo Jun, CF Zhao, Liu Yongming, Saylor Georgia, Athas Grace, Troxclair Dana, Hixson James, Vander Heide Richard, Wang Yue, Van Eyk Jennifer

**Author notes:** Co-senior author (M Herrington David, Hixson James, Vander Heide Richard, Wang Yue, Van Eyk Jennifer). **Corresponding Author Information:** David Herrington, MD, MHS, Dalton McMichael Chair in Cardiovascular Medicine, Department of Internal Medicine Medical Center Boulevard \ Winston-Salem, NC 27157.

## Abstract

The inability to detect premature atherosclerosis significantly hinders implementation of personalized therapy to prevent coronary heart disease. A comprehensive understanding of arterial protein networks and how they change in early atherosclerosis could identify new biomarkers for disease detection and improved therapeutic targets. Here we describe the human arterial proteome and the proteomic features strongly associated with early atherosclerosis based on mass-spectrometry analysis of coronary artery and aortic specimens from 100 autopsied young adults (200 arterial specimens). Convex analysis of mixtures, differential dependent network modeling and bioinformatic analyses defined the composition, network re-wiring and likely regulatory features of the protein networks associated with early atherosclerosis. Among other things the results reveal major differences in mitochondrial protein mass between the coronary artery and distal aorta in both normal and atherosclerotic samples – highlighting the importance of anatomic specificity and dynamic network structures in in the study of arterial proteomics. The publicly available data resource and the description of the analysis pipeline establish a new foundation for understanding the proteomic architecture of atherosclerosis and provide a template for similar investigations of other chronic diseases characterized by multi-cellular tissue phenotypes.

**Highlights:**

- LC MS/MS analysis performed on 200 human aortic or coronary artery samples
- Numerous proteins, networks, and regulatory pathways associated with early atherosclerosis
- Mitochondrial proteins mass and selected metabolic regulatory pathways vary dramatically by disease status and anatomic location
- Publically available data resource and analytic pipeline are provided or described in detail

## Introduction

At the molecular level atherosclerosis can be defined as an assembly of hundreds of intra and extra-cellular proteins that jointly alter cellular processes and produce characteristic remodeling of the local vascular environment. Ultimately, these proteomic changes produce the lesions responsible for most ischemic cardiovascular events. Unfortunately, current methods to treat and prevent cardiovascular disease focus on antecedent risk factors that are not deterministic of these changes, or on anatomic manifestations of disease that are only evident long after these proteomic changes are underway. To improve early disease detection, and to interrupt the disease process before clinical consequences occur, it is necessary to recognize and understand the specific patterns and dynamic features of arterial protein networks that constitute the molecular signatures of healthy and atherosclerotic arterial tissues.

Previous studies have described features of the arterial proteome in murine ^1–3^ and cell models of atherosclerosis ^4,5^ and in limited numbers of human arterial samples with and without atherosclerosis ^6–18^. To date there has not been a comprehensive survey of the human arterial proteome based on a large number of human coronary and aortic samples, using contemporary LC-MS/MS technology and subsequent identification of the proteins that signify presence of pre-clinical atherosclerosis. Accordingly, we established a tissue acquisition, mass-spectrometry analysis and statistical and bioinformatic pipeline to characterize the human coronary and distal aortic arterial proteome and to identify those proteins, networks and pathways most strongly associated with early atherosclerotic lesions. Detailed analyses of the detected proteins reveal several key features of the coronary and aortic proteome in health and disease as outlined below, and demonstrate how the data resource may be used to guide more targeted functional research concerning the proteomics of early atherosclerosis.

## Results

### Global Analysis of Coronary and Abdominal Aortic Proteomes Identifies Novel Arterial Proteins and Scale-Free Network Topologies

Based on stringent quality control and calling criteria a total of 1925 unambiguous protein groups (hereinafter referred to as “proteins”) were identified in one or more left anterior descending (LAD) or distal abdominal aorta (AA) samples, including 974 proteins present in 50% or more of the LAD or AA samples (**Supplementary Table 1**). The 974 proteins represent a wide range of biological processes, molecular functions, cellular components and canonical pathways^19^ (**Supplemental Figs. 1a-c and Supplemental Tables 2-5**). Of the 1925 proteins, 274 have not been previously described in twelve prior studies that directly analyzed protein content of human arterial tissues (**Supplemental Table 1**); including 52 proteins present in more than 50% of either the LAD or AA samples. (All but one of these 52 proteins were, nevertheless, predicted based on RNASeq analysis of human coronary and aortic transcriptional databases (GTEX)).

The 944 unique proteins identified in the LAD exhibited a scale-free network topology typically seen in complex adaptive networks of molecular or cellular constituents from a variety of living organisms ^20–22^ (**Fig. 1**). Furthermore, these proteins included several distinct and reproducible co-expression modules that roughly correlated with specific cellular functions and locations such as mitochondrial proteins involved with cellular respiration (Red module), nuclear proteins involved with chromatin assembly and organization (Turquoise module), and extra cellular matrix proteins (Brown module). A similar scale-free topology and functional modular structure was also evident in the protein data from the AA (**Supplemental Fig. 2.**) A scale-free topology suggests that the arterial proteome may arise from a complex adaptive system with properties such as self-organized criticality, emergence, and resilience.^21,23,24^ Complex adaptive systems such as this are uniquely well-suited for description using graph theory and network or non-linear dynamic modelling to reveal functional insights that may be less evident from more conventional linear conceptual models and methods ^25–27^

**Figure 1.**
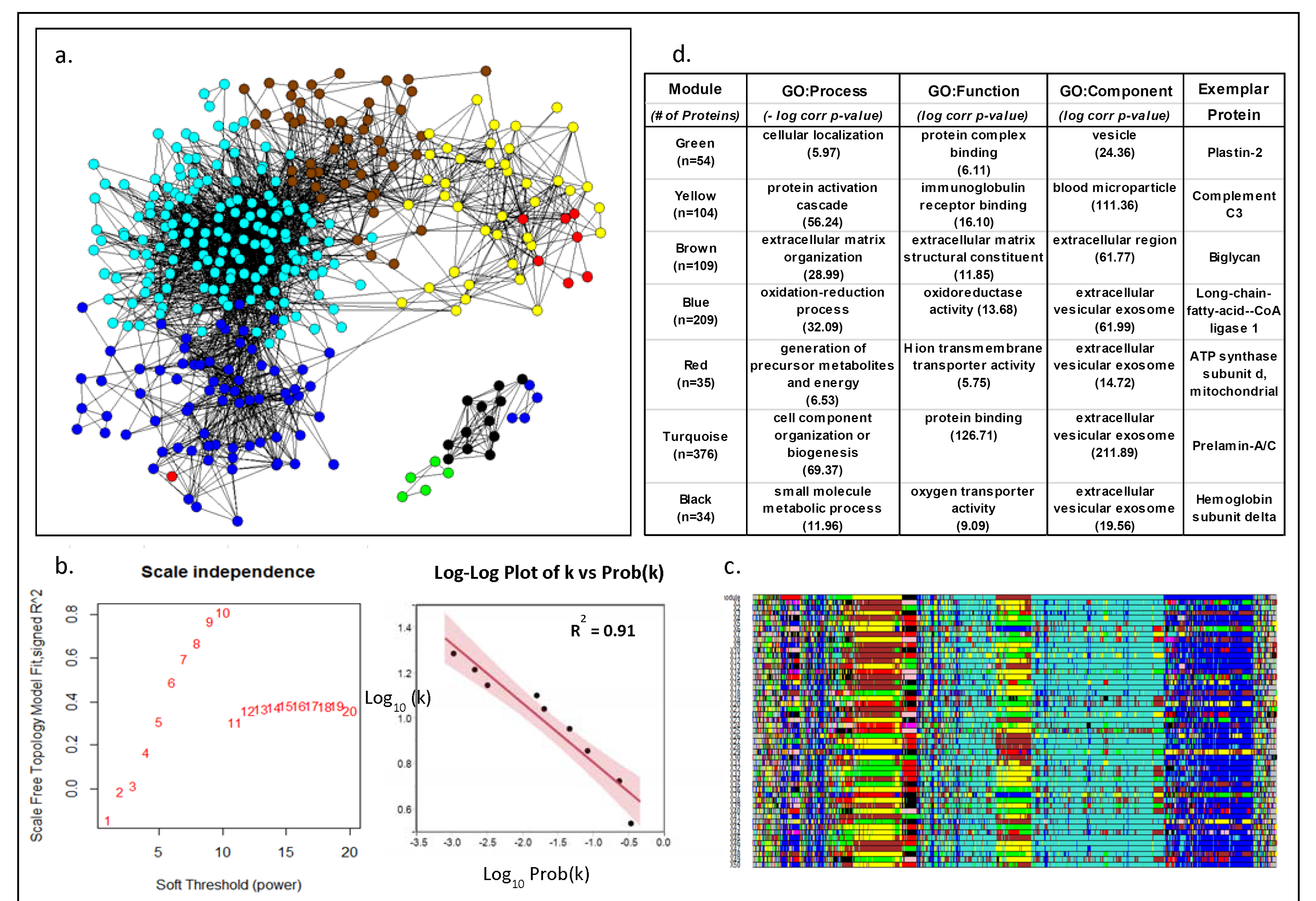
Weighted Co-Expression Network Analysis of Human LAD Proteins. **a.** Adjacency map of LAD proteins color coded by module assignment based on hierarchical clustering of the topological overlap matrix (TOM)-based dissimilarity measure. For clarity of presentation only nodes (proteins) with at least one edge (adjacency measure (k)) > 97.5%tile are shown, b. Left panel shows that scale-free topology is best approximated when the adjacency power parameter (β= 10. Right panel shows the log-log plot of adjacency (k) vs prob(k) with (β= 10, confirming the power-law relationship in the connectivity of the expressed proteins, c. Module assignment for fifty 90% random samples of the data illustrating the overall stability of the modular structure of the protein expression patterns. Colors are assigned according cluster size which may vary with each random sample. As a result actual color assignment may vary from run to run, but module membership remains relatively stable, d. Top non-redundant GO Terms with Bonferroni corrected p-values and an exemplar protein for each module.

### Comparison of Normal Coronary and Aortic Proteomes Reveal Significant Differences in Mitochondrial Protein Mass

Several hundred proteins were detected in the LAD but not in the AA samples (**Supplemental Figure 3**). This pattern was observed when limiting the analysis to completely normal samples (n=30 in each territory) or when limiting the proteins to those present in > 50% of the LAD and/or AA samples. GO term analysis of the proteins exclusively detected in the LAD indicated significant enrichment of mitochondrial proteins (p-value range 1.7×10^−6^ to 1.8×10^−28^).

To confirm this apparent differential abundance of mitochondrial proteins between LAD and AA samples we performed a more sensitive data independent acquisition MS (SWATH) analysis of entirely normal samples (n=30 in each location), focusing on n=114 mitochondrial proteins involved with fatty acid metabolism, oxidative phosphorylation, TCA cycle and mitochondrial biogenesis. To account for possible site and sample differences in cellular material or protein extraction yields the quantitative results were adjusted for several housekeeping proteins, a smooth muscle cell specific marker protein and age and sex of the autopsied cases. Overall, mitochondrial proteins were 1.98-fold more abundant in the LAD compared with the AA (p<0.001) including a 2.25 fold increase in oxidative phosphorylation proteins (p<0.001) and an isolated >10-fold excess of inorganic pyrophosphatase (p<0.001; **Figure 2**). A similar comparison of extracellular matrix proteins revealed only a small, albeit statistically significant (p=0.01) 6% excess in ECM proteins in the AA samples compared with the LAD (**Supplemental Figure 4**). (A notable exception was tenascin which had a >10-fold excess in AA samples compared with LAD (p<0.0001).)

**Figure 2.**
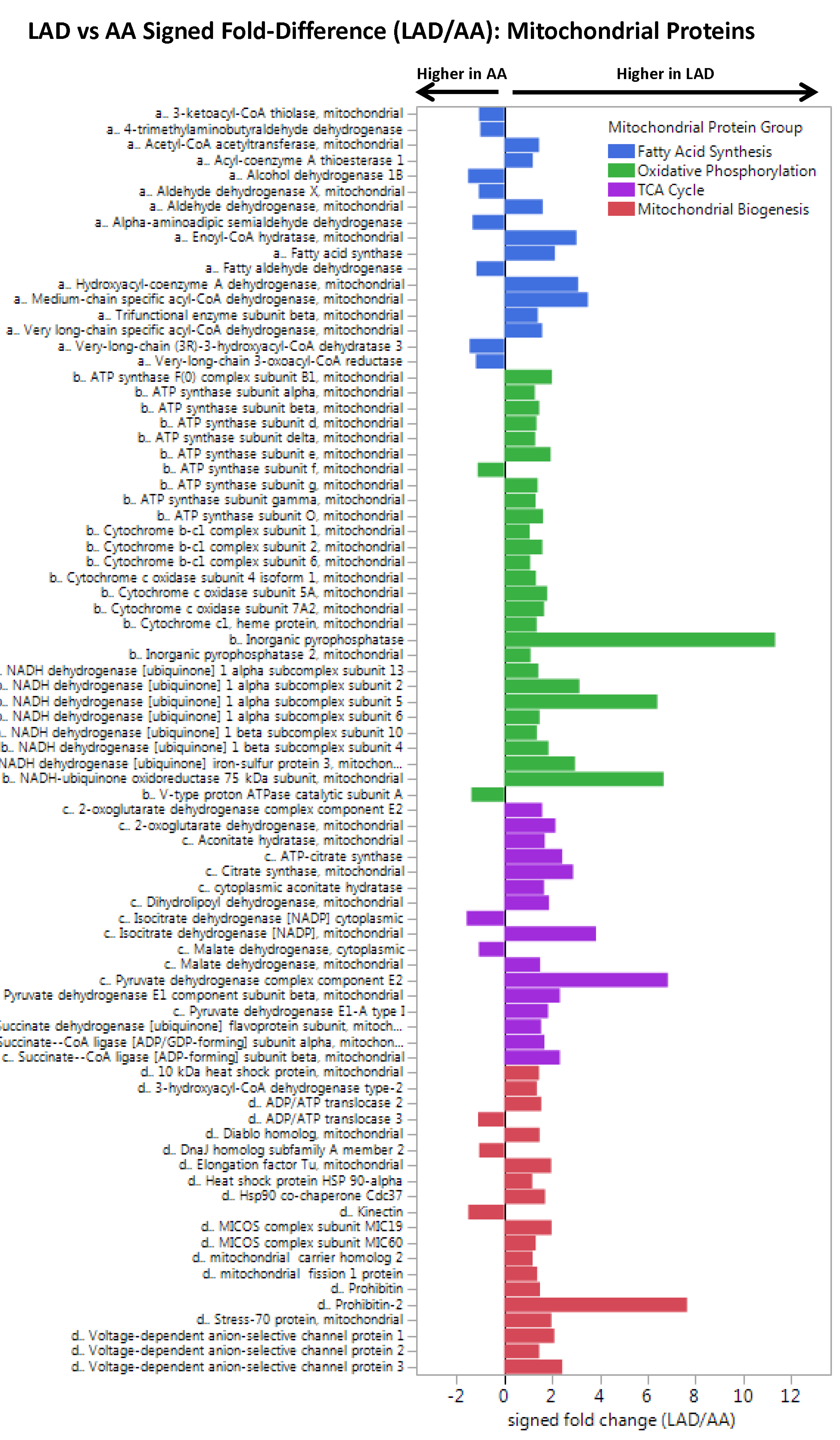
Comparison of Mitochondrial Proteins in Normal LAD and AA Samples. Data independent MS (SWATH) analysis of completely normal LAD and AA samples (n=30 from each anatomic location) with adjustment for age, sex, MYH11, RABA7A, TERA, G6PI. LAD vs AA MANOVA p-values by proteins class: fatty acid metabolism, p = 0.04, oxidative phosphorylation p < 0.0001, TCA p < 0.0001, mitochondrial biogenesis p < 0.0001.

These data suggest fundamental differences in mitochondrial mass and potential aerobic capacity between LAD and AA tissues, possibly reflecting the differing energy requirements of these two arterial tissue types.

This is exemplified by the dramatic excess of inorganic pyrophasphatase in the LAD compared to the aorta suggesting enhanced capacity to initiate fatty acid oxidation. This heterogeneity in the proteomic profile of two arterial tissues emphasize the need for arterial anatomic specificity when characterizing the proteomics and functional biology of arterial tissues – especially if considering mechanisms or interventions that involve metabolic pathways (see below)

### Atherosclerotic tissues in both the LAD and AA present a proteomic profile consistent with TNF activation; however, the LAD also provides evidence of inhibition of PPAR-a, PPAR-y, and insulin receptor regulated proteins, a pattern that is not evident in the AA

To define the proteomic profile of early atherosclerosis two complementary phenotyping strategies were used. First, each sample was graded by a vascular histopathologist according to %surface area involvement of normal intima (NL), fatty streak (FS) or fibrous plaque (FP). Extensive regression analyses (MANOVA, GLM, Ordinal Regression, Elastic Net) were performed to identify proteins individually or jointly associated with %FP in the LAD and AA samples (244:106 **Supplemental Tables 6–7, Supplemental Figure 5**). The internal validity of these data is evidenced by the presence and consistency across anatomic locations of several known protein markers of atherosclerosis from model systems among the top hits (e.g. Apo B-100, Ig mu chain C, CD5 antigen-like, Plastin-2, Tenascin, Thrombospondin-1, Cathepsin B, and Vitronectin.) ^1^

Separately, convex analysis of mixtures^22^ (CAM) was used to de-convolve the global protein profiles from individual samples into data-derived tissue phenotypes and to identify marker proteins associated with each data-derived tissue phenotype (**LAD: Figure 3, AA: Supplemental Figure 6**). This approach has the advantage of incorporating information about tissue phenotype from the global protein profiles that is independent of the pathologist visual inspection of the arterial samples. The data-derived tissue phenotypes, and associated FP marker proteins roughly correlated with results based on pathologic assessment of extent of disease, but retained sufficient variation from pathology based results to suggest complementary information was present (**Supplemental Figure 7**).

**Figure 3.**
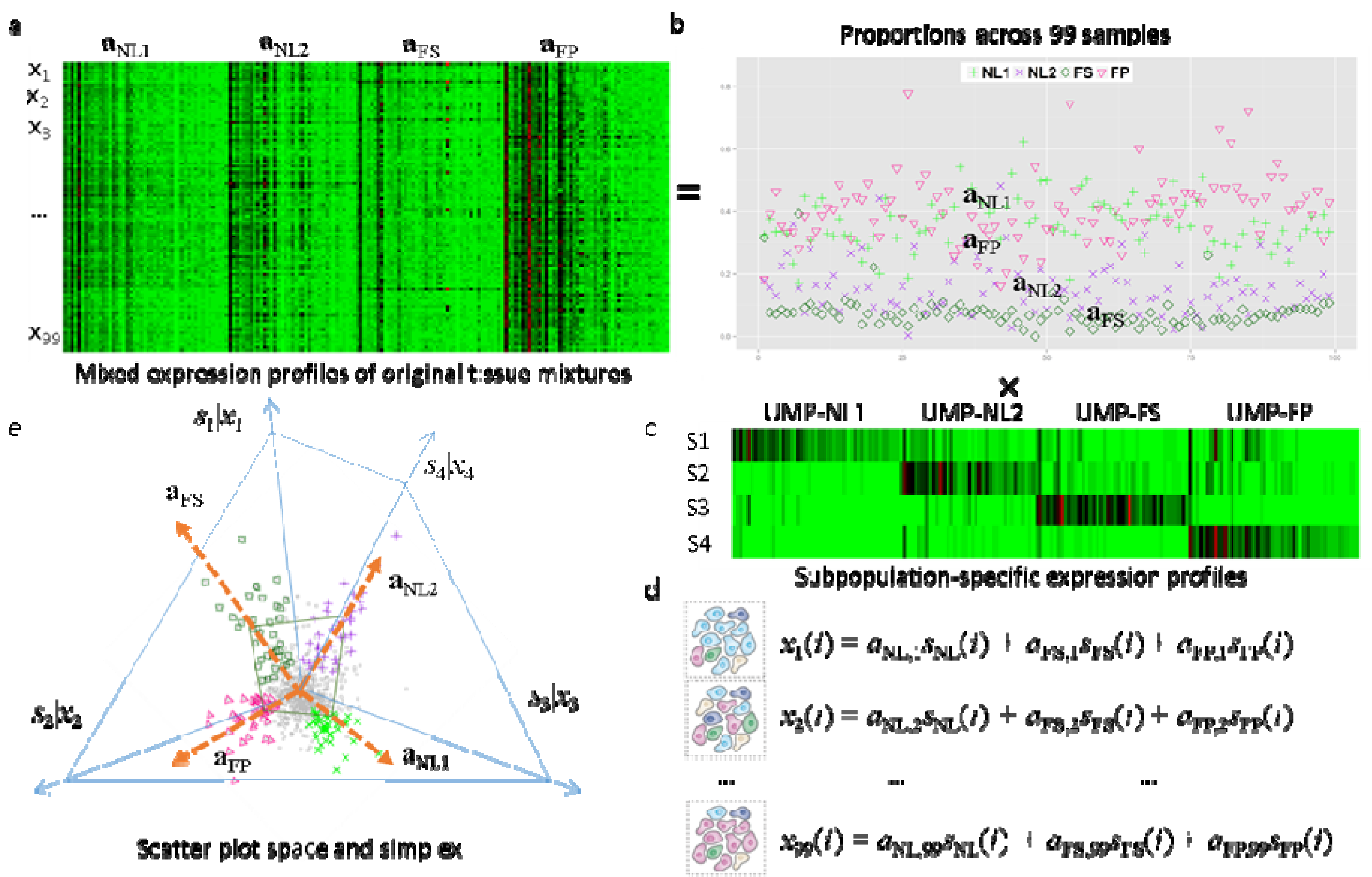
Ccnvex Analysis of Mixtures of LAD Protein Data. a. Heatmap 3^f^ mixed expressions of UMPs in 99 LAD samples. b. Estimated proportions of NL1, NL2, FS, and FP across 99 LAD samples. c. Heatmap of subpopulatlon-specific expressions of UMPs.d. Mathematical descrition on the protein expression readout of multiple distinct subpopulatlons. e. Geometry of the mixing operation In scatter space that produces a compressed and rotated scatter simplex whose vertices host subpopulatlon-speclflcUMPs and correspond to mixing proportions.

To take full advantage of both phenotyping strategies we used principal component analysis and hierarchical clustering of the pathologist-and CAM-derived phenotype data to produce a patho-proteomic classification for each LAD sample (**Fig. 4**). The clusters at the extremes of the first principle component identified samples highly enriched with FP or NL tissue (FP: n=15; NL: n=30) with little or no confounding from fatty streaks from either a gross pathology or global proteomic perspective. Comparing these FP and NL enriched samples identified eighty-nine (n=89) individual proteins with ≥ (+/-) 1.7 fold-difference and a t-test q-value of ≤0.05 for FP vs NL (**Supplemental Tables 8 and 9**). Bioinformatic functional analysis of these atherosclerosis associated proteins revealed a pattern consistent with activation of the TNF-α pathway (p=2.64E-07), but also inhibition of insulin receptor, PPAR-± and PPAR-γ pathways (p=4.22E-10, 2.42E-13, 8.56E-16 respectively) (**Fig. 4, Supplemental Table 10**). A similar analysis of the atherosclerosis proteins in the AA samples (FP: n= 9, NL: N= 18) confirmed a core group of n=19 early atherosclerosis-associated proteins that were shared across both anatomic territories (**Supplemental Table 11–13**) and also produced a pattern consistent with TNF activation similar to the LAD (p=1.92E-05). However, in the AA sample proteomes there was no evidence of inhibition of the insulin receptor, PPAR-α or PPAR-γ pathways (**Supplemental Tables 10**).

**Figure 4.**
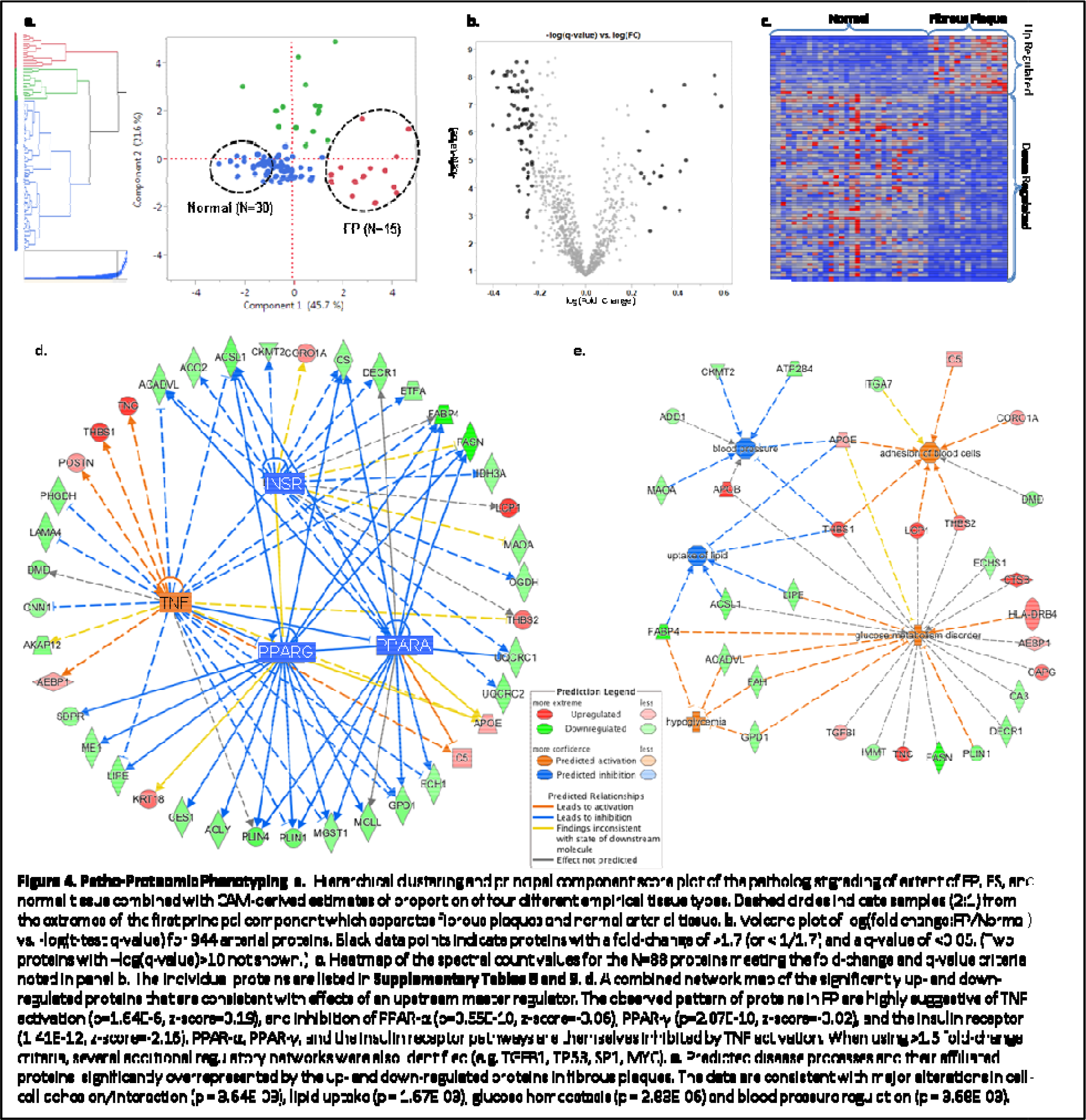

### Differential network analysis of coronary and aortic proteomes indicate divergent mitochondrial dynamics in the setting of atherosclerosis characterized by reduced mitochondrial mass in coronary arteries that is not evident in in the distal abdominal aorta

An important feature of complex adaptive systems is the potential for network topologies to change under different conditions. Accordingly, we used differential dependent network (DDN) analysis of the FP-associated proteins identified above to select proteins that were pivotal in the re-wiring of the network structure between NL and FP in the LAD samples (**Figure 5**). Analysis of n=26 re-wiring hub proteins revealed significant enrichment of TCA proteins (p=4.8×10^−6^). Subsequent analysis of individual TCA proteins documented an average 60% reduction in TCA proteins in FP enriched samples vs. NL.

**Figure 5.**
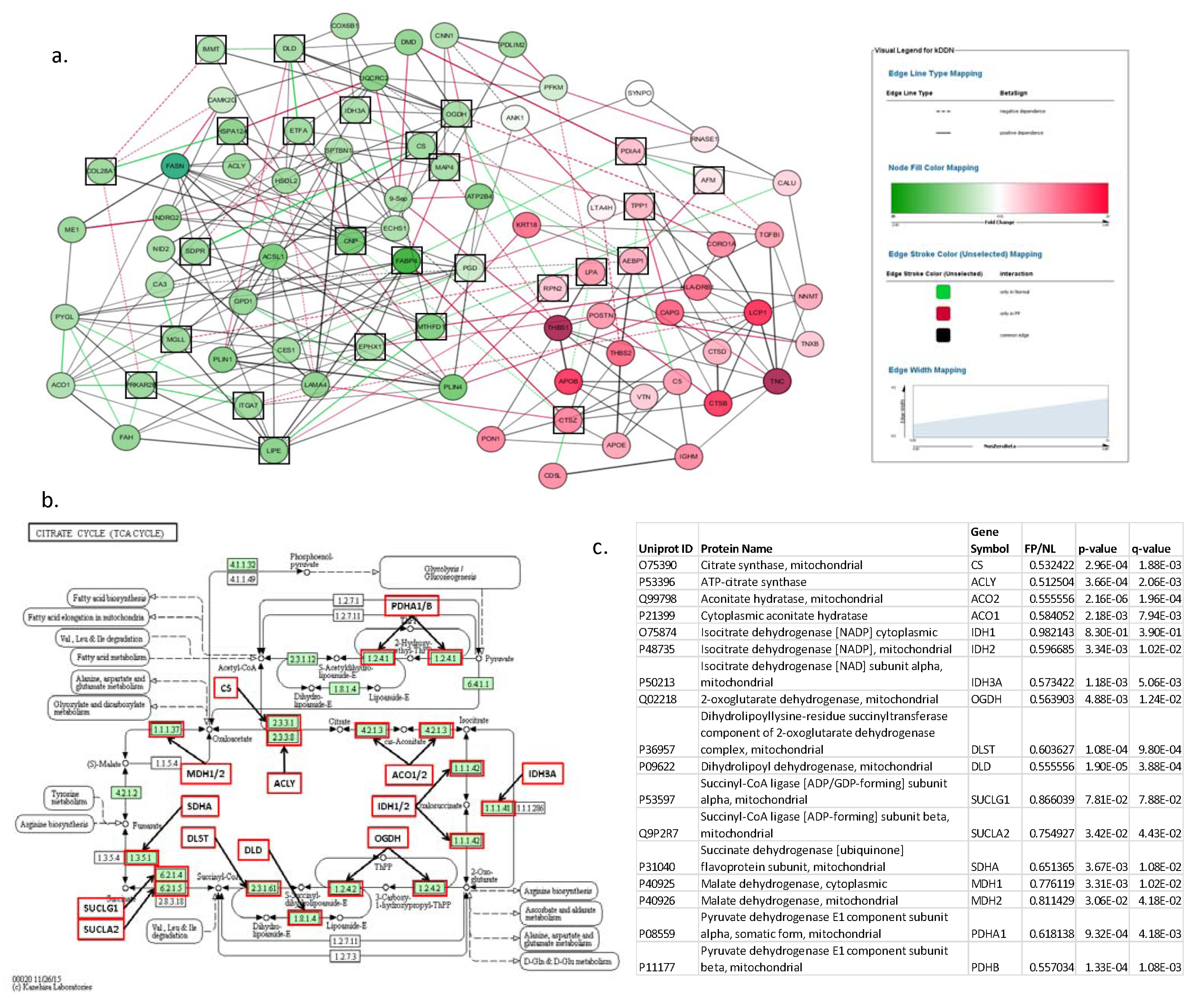
Differential Network Analysis of FP Associated Proteins in the LAD. **a.** Plot depicts the re-wiring of protein networks between FP and NL samples. Green nodes are up-regulated and red nodes are down-regulated in FP samples. Green edges indicate significant correlation in normal samples not present in FP samples. Red edges indicate significant correlation in FP samples not present in normal samples. Black squares indicate differential network “hub proteins” (ie. proteins with different couplings to network partners in both FP and NL samples). Go term analysis of the differential network hub proteins indicated significant enrichment of TCA proteins (p=4.8×l0-6). **b.** TCA cycle proteins with MS data available for additional analysis. Every protein indicated by a red box was quantitatively lower in FP samples after adjustment for housekeeping proteins, age and sex. **c.** Statistical comparison of TCA proteins in FP vs NL LAD samples.

To confirm and extend this observation, SWATH analysis of a broad-based panel of n=114 mitochondrial proteins was performed comparing FP vs NL in the LAD (**Figure 6**). The results documented a consistent reduction of a wide range of mitochondrial proteins in the FP samples compared to NL samples after adjustment for housekeeping and vascular smooth muscle cell marker proteins, age and sex. In contrast, a similar analysis of the same proteins in AA samples revealed a much less consistent and non-statistically significant pattern of mitochondrial protein suppression. To determine if this was a mitochondrial specific phenomenon we also performed targeted SWATH analysis of a panel of ECM proteins (n=77) and found a modest atherosclerosis-associated increase of ECM proteins in both territories (LAD mean fold-increase = 1.25, MANOVA p-value= 0.02; AA mean fold increase = 1.78 fold increase, p-value = 0.017); although there were specific examples of anatomic discordance that deserve further study (e.g. laminins and nidogens) (**Supplemental Figure 8**).

**Figure 6.**
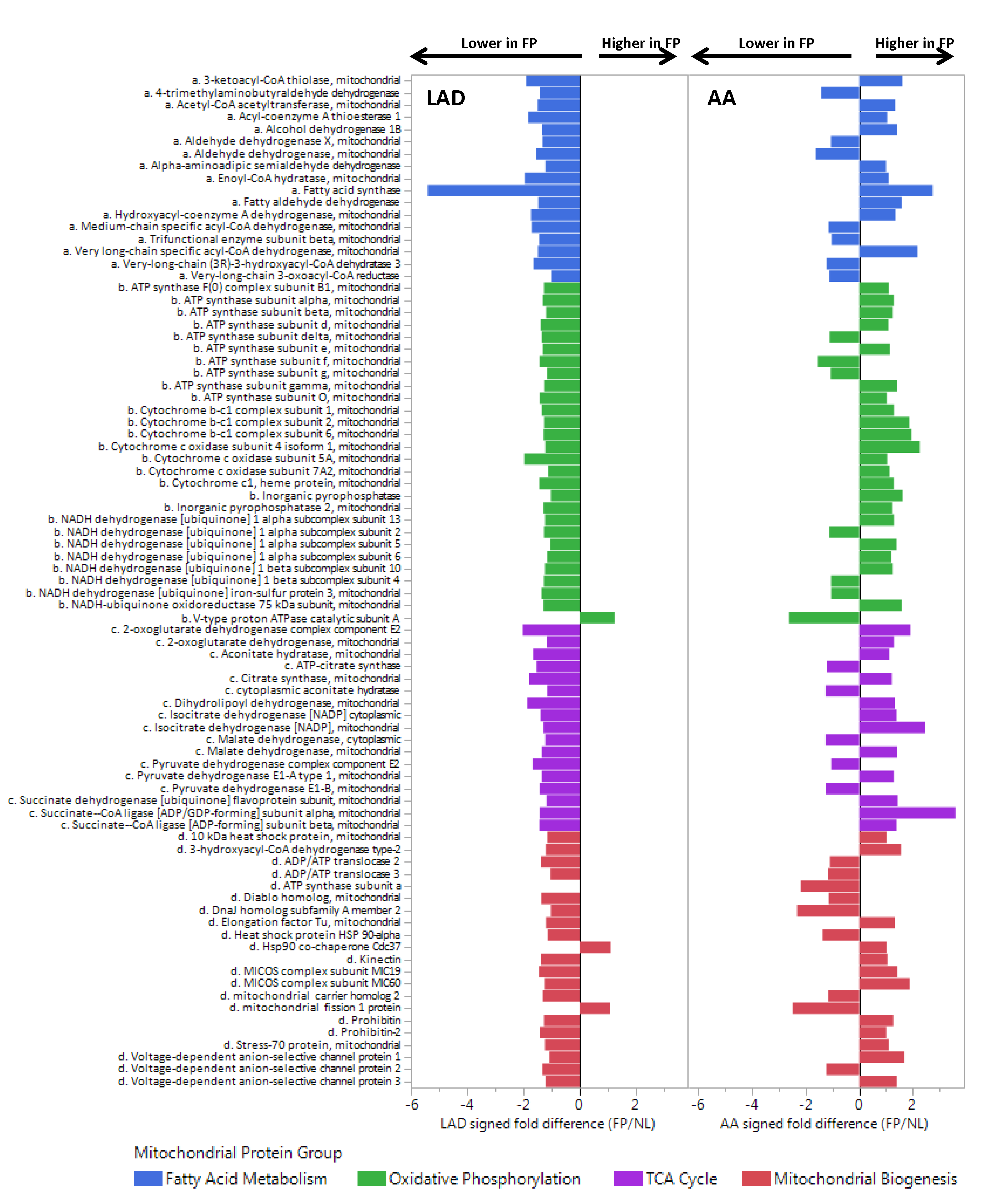
Comparison of Mitochondrial Proteins in FP vs NL Samples from the LAD and AA. Data independent acquisition (SWATH) analysis was used to compare a targeted set of mitochondrial proteins in n=15 FP and n=30 NL LAD samples after adjustment for age, sex, MYH11, RABA7A, TERA, G6PI. Histogram bars indicate relative difference between FP and NL samples in each anatomic location. LAD MANOVA p-value for each mitochondrial protein group: p <0.0001 for each group; AA MANOVA p-values for each mitochondrial protein group: p = n.s. for each group.

## Discussion

The results here provide a comprehensive survey of human coronary and aortic proteins and identify individual proteins, protein networks and regulatory pathways indicative of early atherosclerosis. These data can be distinguished from prior work because of the number and diversity of proteins identified, the fact that these proteins were obtained from a large number of human coronary and aortic arterial samples where the effects of local tissue context, natural variation in human subjects, and a range of arterial tissue phenotypes from normal to early atherosclerosis are manifest. Several analytic methods were used to illuminate complex networks involving many proteins that jointly signify arterial health and disease. Importantly, the publicly accessible data generated by this work, and the analytic methods depicted here represent foundational resources that could lead to a wide range of future research concerning human arterial protein biology.

There are several notable findings among the many results reported here. First, the human arterial proteome exhibits features consistent with a complex adaptive network, reiterating the concept that complex adaptive networks are an overarching organizational feature common to many biologic systems^28^. Looking at the arterial proteome through the lens of complexity theory including concepts such a scale-invariance (as documented here), self-organized criticality, emergence, and resiliency ^21,23,24^ may produce insights that have escaped more narrowly focused investigations of a smaller number of proteins or a more linear and deterministic framework. The differential dependent networks presented here emphasize this point by defining dynamic features of complex protein networks that signify normal versus atherosclerotic arteries and using key elements of these dynamic networks to discover important aspects of the proteome in atherosclerosis that were not initially evident by other means.

Second, the data document significant anatomic variation in the abundance of mitochondrial proteins in normal arterial samples, suggesting that coronary arteries likely have considerably greater aerobic capacity than the distal aorta. This fits our understanding of the different embryology^28^ and normal physiologic roles of these distinct regions of the arterial system with the coronary arteries having greater energy requirements associated with regulation of coronary blood flow compared to the distal aorta which serves primarily as passive conduit for blood delivery to the lower extremities. To our knowledge this substantial variation in mitochondrial protein mass between coronary and aortic tissues in humans has not been previously documented.

Third, the data indicate profound anatomic differences in key metabolic regulatory pathways and mitochondrial dynamics in the setting of atherosclerosis. Specifically, the data reveal a broad-based reduction of mitochondrial protein mass and a proteomic pattern consistent with inhibition of PPAR-α, PPAR-γ, and insulin receptor regulated pathways in atherosclerotic coronary arteries, but not in similarly diseased distal abdominal aortic samples. These provide critical anatomic specificity to the growing body of evidence that mitochondrial dysfunction and altered mitochondrial dynamics are also central features of atherosclerosis ^29 15 30^, and may provide a mechanistic framework to explain the anatomic dimorphism between coronary and peripheral arterial disease with respect to conventional and genetic risk factors for atherosclerosis ^31,32^ It remains to be determined whether the reduction in mitochondrial protein mass in coronary atherosclerosis is a consequence of oxidative, or Ca^++^-mediated mitochondrial damage and mitophagy or impaired mitochondrial biogenesis, or both. It is interesting to note that the concurrent increases in the lysosomal proteases cathepsin B, D and Z in the atherosclerotic samples may signify enhanced lysosomal biogenesis useful for targeted degradation of damaged mitochondria ^29^.

The data also reassuringly highlight numerous individual proteins and a global proteomic pattern of TNF activation in the FP samples consistent with several decades of research documenting the role of inflammation in the pathogenesis of atherosclerosis. ^33^ However, even the more familiar indicators of TNF activation and inflammation exhibited substantial anatomic variation from LAD to distal aorta in our data, an observation that was made possible by the extended proteomic depth of coverage in the current study.

Recognizing the anatomic heterogeneity with respect to metabolic enzyme profiles and mitochondrial function has important implications for research on arterial health and disease. Many of the pathogenic mechanisms and targeted therapies for atherosclerosis are directly related to fatty acid metabolism or aerobic energy biosynthesis and redox homeostasis ^34^. Unfortunately, a great deal of our understanding of arterial biology has been generated without regard to anatomic distribution. For instance many animal models of atherosclerosis rely on aortic, femoral or carotid manifestations of disease with the assumption that findings are generalizable to human coronary arteries. Likewise the overwhelming majority of contemporary human arterial proteomic data have been generated from carotid arteries – typically severely diseased endarterectomy specimens. The current data suggest that a complete understanding of the pathogenesis and prevention of human coronary artery atherosclerosis, especially as it relates to metabolic mechanisms and mitochondrial function, may require more anatomically specific methods of interrogation. It is also interesting to consider whether anatomic variations in the arterial proteome may be illuminating about other forms of arterial disease that have distinct anatomic distributional features (e.g. Marfan’s syndrome, Kawasaki disease, fibromuscular dysplasia, etc).

Several limitations in the current work should be considered. First, although a comprehensive analysis of 200 human arterial specimens is a considerable technical accomplishment resulting in much greater power to discern real patterns in the data, and despite the fact that we used stringent statistical methods and other information theoretical considerations to minimize false positives (generally to < 5%), there remains the possibility of residual confounding or limited statistical power which could obscure both positive and negative associations. This is especially true for the AA samples where the number of significantly diseased arterial samples was more limited than in the LAD. The fact that many of the sentinel findings generated by the initial data dependent MS approach were subsequently confirmed and strengthened by a separate data independent acquisition MS method (which typically utilizes different peptides for protein identification) provides further support for the validity of the findings. Second, the fact that the samples were all collected post-mortem raises questions about possible effects of death on the stability of individual proteins or specific cellular functional processes. If such effects are differentially manifest in normal versus atherosclerotic tissues, this could give the appearance of differences by disease status that are not present during life. It is reassuring that many of the individual proteins identified here overlap with findings from animal models of atherosclerosis and *in-vitro* studies where the effects of post-mortem hypoxia and cessation of cellular metabolic activity are more easily minimized. Third, the observed differences in the proteome between normal and diseased samples represent a unique, albeit macroscopic view of the proteomic architecture of human coronary and distal aortic atherosclerosis based on analysis of hundreds of proteins. Much more work is required to clarify the functional role of the specific proteins, protein networks and pathways identified here, and to establish if there is therapeutic value in manipulating them – perhaps even in an anatomically specific manner.

In summary, these data represent the most comprehensive description of the human coronary and aortic proteome to date and reveal numerous proteins, networks and pathways that are strongly indicative of early atherosclerosis. They also indicate fundamental differences in mitochondrial dynamics between the coronary artery and the distal aorta in both normal and atherosclerotic conditions. The data analyses highlight the value of new methods to extricate tissue phenotypes from heterogeneous tissue samples and depict dynamic features of protein networks that vary as a function of disease state and anatomic location. These data and methods establish a new foundation for future research to better understand the human arterial proteomic architecture in health and disease.

## Author Contributions

D.H., J.V.E., Yue W., R.V.H., J.H. designed the project, supervised the execution of the research plan, and made significant contributions to the interpretation and presentation of the results; C.M. and S.P. provided critical project management and scientific direction concerning specific aspects of the data acquisition and bioinformatic analyses; Z.F. and V.V. were primarily responsible for generation and annotation of the mass-spectrometry data; G.Y, L.C, Y.F., Y.W., J.G., G.S. performed data analysis; T.H. and Y.L. contributed to genomic analyses and interpretation; D.T. and G.A. and were responsible for collecting, grading, storing and shipping the pathologic specimens.

## Acknowledgments

This work was supported by 1R01HL111362 from the, National Heart, Lung and Blood Institute of the National Institutes of Health

## Methods

### Arterial Sample Acquisition and Pathology Grading Methods

Male and female cases of any race, aged 18-50 years (men) or 18-60 years (women) with no ante mortem clinical suspicion of coronary disease autopsied < 24 hours of death were eligible for inclusion. This report includes data from the first 100 adult coroner cases included in the study (age range:15-55 yrs., 75% males, 67% White, 26% Black, 7% Other). The Medico-Legal Death Investigators obtain signed family consent for retrieval of anatomic specimens prior to the autopsy. During the autopsy the pathologist dissects the aorta (ligamentum arteriosum to aortic bifurcation) and LAD and removes branching arteries and adventitial or epicardial adipose tissue. Both the aorta and LAD are opened longitudinally, cleansed of blood and photographed (**Supplemental Fig. 9a and b**). In each of the regions to be sampled, the pathologist inspects the intimal surface and categorizes the type and percent involvement of the following atherosclerotic changes: a) fatty streaks (FS); b) fibrous plaques (FP); c) complicated lesions (CL); and d) calcified lesions (CO), and records the data on a data collection form. Consistent with the age and cause of death (trauma), the fibrous plaques were almost exclusively early lesions-only one specimen had any macroscopic evidence of calcification and <3% of samples had any evidence of plaque hemorrhage, ulceration or thrombosis. Next the pathologist collects up to 1 gram of tissue from each of three standardized sections of the thoracic and abdominal aorta and two sections from the coronary artery ^35^ (**Supplemental Fig. 9c and d**). The specimens are snap frozen in cryotubes with liquid nitrogen and remain in a 34L VWR Cryogenic Dewar until transferred to a −80 freezer equipped with temperature alarms and automated generator back-up systems in the Department of Pathology at LSU. Additionally, targeted “pure” samples of grossly normal (non-lesion) and, if available, grossly atherosclerotic (lesion) are taken and divided into three 100mg portions and 1.) snap frozen, 2.) placed in RNALater solution and frozen, and 3.) placed in formalin and stored at room temperature for possible future immunohistochemistry analyses. A 50 gram sample of liver is also collected and frozen for future studies. All samples received at LSU are checked for label accuracy and entered in a database using an unlinked anonymous code for further processing, analyses and storage.

### Protein Extraction, MS Analysis and Protein Identification

Aorta and LAD tissues were pulverized in liquid nitrogen and homogenized in 8M urea, 2M thiourea, 4% CHAPs and 1% DTT using Dounce homogenizer for 100 strokes, centrifuged at 16000 rpm for 20 mins. Protein concentration of the supernatant was assessed by CB-X assay kit (G-Biosciences MO, USA). 100 μg of protein was precipitated using 2-D clean-up kit (GE Healthcare MA, USA) and then reconstitute in 6M urea, 50mM ammonium bicarbonate. The protein was reduced, alkylated and digested with trypsin (1:20). Peptides were desalted using Oasis HLB 96-well Plate (Waters MA, USA). A total of 2.0 μg of peptides per sample were then analyzed using label-free quantification on a reversed-phase liquid chromatography tandem mass spectrometry (RPLC-MS/MS) online with an Orbitrap Elite mass spectrometer (Thermo Scientific, USA) coupled to an Easy-nLC 1000 system (Thermo Scientific, USA). Peptides concentrated on a C18 trap column (Acclaim PepMap 100, 300 μm × 5 mm, C18, 5 μm, 100 Á: maximum pressure 800bar) in 0.1% TFA, then separated on a C18 analytical column (Acclaim PepMap RSLC, 75 μm * 15 cm, nano Viper, C18, 2 μm, 100 Á) using a linear gradient from 5% to 35% solvent B over 155 mins (solvent A: 0.1% aqueous formic acid and solvent B: 0.1% formic acid in acetonitrile; flow rate 350 nL/min; column oven temperature 45 °C). The analysis was operated in a data-dependent mode with full scan MS spectra acquired at a resolution of 60,000 in the Orbitrap analyzer, followed by tandem mass spectra of the 20 most abundant peaks in the linear ion trap after peptide fragmentation by collision-induced dissociation (CID).

To eliminate batch effect, the same parameters were used for mass spectra acquisition and the peptides from each individual were analyzed randomly in one batch. To minimize cross contamination, a blank run was performed between each sample. To monitor column performance, 200fmol BSA digested peptides were analyzed to make sure elution time of same peptide were within 0.2 minutes and signal intensity and total spectra counts variation were less than 10%. One LAD and one AA sample (from different subjects) were excluded because of poor protein yield leaving n=99 samples from each territory for analysis.

The MS/MS data obtained from the Orbitrap Elite were converted to mzXML and mgf format using Msconvert version 3.0.3858 from ProteoWizard ^36^ for peaklist generation. All data were searched using the X!Tandem ^37^ algorithm version 2009.10.01.1 and OMSSA ^38^ algorithm version 2.1.9. The dataset was searched against the concatenated target/decoy ^39^ Human Uniprot ^40^ database as of July 24, 2015, with only reviewed and canonical sequences used. The search parameters were as follows: Fixed modification of Carbamidomethyl (C) and variable modifications of Oxidation (M), Phosphorylation (STY); Enzyme: Trypsin with 2 maximum missed cleavages; Parent Tolerance: 0.050.08 Da; Fragment tolerance: 1.00 Da. Post-search analysis was performed using Trans Proteomic Pipeline ^41^ version v4.6, rev 1 with protein group and peptide probability thresholds set to 90% and 90%, respectively, and one or more distinct peptide required for identification. PeptideProphet ^42^ was used for peptide validation from each individual search algorithm and iProphet ^43^ was used to merge results from the separate algorithms and further refine the identification probabilities. Lastly, ProteinProphet ^44^ was then used to infer protein identifications from the resulting combined peptide list and perform grouping of ambiguous hits. Protein Group and Peptide False Discovery Rates were calculated automatically using a target-decoy method for the above probability thresholds (0.72% and 0.06% respectively). Protein isoforms were only reported if a peptide comprising an amino acid sequence that was unique to the isoform was identified. Label-free quantification of each protein was performed using weighted spectral counting ^45^. The distribution of protein spectral counts for each data file was inspected to verify equivalence in sample loading and instrument performance between files. Based on this analysis, a median normalization (adjustment of the raw spectral counts for each protein of a given file to the median of all spectral counts observed for the same file) was performed (**Supplemental Fig. 10**).

The final non-redundant protein groups were analyzed through the use of IPA (Ingenuity® Systems, www.ingenuity.com) to generate network, functional and pathway analyses. The MS/MS proteomics data have been deposited to the ProteomeXchange Consortium (http://www.proteomexchange.org) via the PRIDE partner repository ^46^ with the dataset identifier (# to be added at the time of publication).

### Post-processing Quality Control and Data Imputation

A total of 1925 unambiguous proteins were detected in one or more of the 99 LAD samples. Of these proteins 944 had ≤50% missingness and 375 had no missingness in all 99 samples (**Supplemental Fig. 11a, d**). PCA analysis of the 375 proteins with complete data from every sample failed to identify significant sample outliers or important batch effects (**Supplemental Fig. 11c**). Missing values for the 944 proteins with ≤ 50% missingness were imputed using a low rank approximation derived from non-linear iterative partial least squares (NIPALS) PCA ^47^ (**Supplemental Fig. 11b**). All subsequent analyses including regression, WCNA, and CAM were performed on this imputed data set. Exploratory regression analysis of proteins with > 50% missingness using missingness encoding failed to identify proteins with compelling association with disease and these proteins were subsequently dropped from further consideration. In a similar manner the AA samples revealed 1495 unambiguous proteins including 725 with <50% missingness and no important batch effects or extreme outliers.

### Analysis of Association with Extent of Atherosclerosis

Three orthogonal strategies (regression, weighted co-expression analysis, convex analysis of mixtures) were employed to identify proteins individually or jointly associated with extent of disease.

#### Regression Models

MANOVA models with adjustment for age, sex, and race were used to model the joint distributions of % intimal surface demonstrating fibrous plaque (FP), streaks (FS), and normal (NL) intima as a function of each individual protein. To account for possible differences in tissue sample volumes, cellular composition or protein yields the models were also adjusted for several housekeeping proteins (Proteasome subunit beta type-2, Small nuclear ribonucleoprotein Sm D3, Receptor expression-enhancing protein 5, Ras-related protein Rab-7a) and Myosin-11 (a vascular smooth muscle cell marker protein). Separately, generalized linear models (GLM) were used to examine the association between individual proteins and %FP or %NL after adjustment for age, sex, race and the same set of housekeeping and VSMC genes. In the models of %FP the reference was, by definition, the % intimal surface demonstrating fatty streaks or normal intima. In contrast, in models of %NL the reference was, by definition, the % intimal surface demonstrating fatty streaks or fibrous plaques. Accordingly we developed separate models using each trait (%FP or %NL) as the dependent variable with the expectation that direction of effect would be inversely related depending on the dependent variable used. Proteins with beta coefficients in the same direction for both %FP and %NL were excluded from further consideration. In a similar manner ordinal regression was also used to examine the association between individual proteins and % FP or % NL treated as three level ordinal variables (FP: 0%, 1¬59%, ≥60%; NL: 100%, 99-61%, <60%). Exploratory models using robust GLM with logit link produced qualitatively similar lists of significantly associated proteins (data not shown).

#### Weighted Co-Expression Network Modelling

The WGCNA package in R was used to identify distinct protein modules among the 94 proteins used for analysis ^48^. A weighted power adjacency matrix was constructed using signed bi-correlations between proteins and mapping the results onto the 0-1 interval. The power parameter was selected such that the topological overlap connectivity (k) of the entire network approximated a scale-free topology. The topological overlap was used to create a dissimilarity matrix for hierarchical clustering to identify modules. To assess stability of module assignments, 50 bootstrap samples of the data were created and each one interrogated using identical WCGNA parameters. Once the modules were determined, regression models were developed to examine the association between module eigengenes and the arterial phenotypes (%FP and % NL) after adjustment for age, sex, and race.

#### Convex Analysis of Mixtures and Protein Expression Differences in Complex Tissues Analysis

Tissue heterogeneity is present in our samples where multiple tissue types are variably mixed and co-exist ^49,50^, representing a major confounder when studying tissue-specific disease markers ^51–53^. We have recently devised CAM, a fully unsupervised data deconvolution method that exploits the strong parallelism between a latent variable model and the theory of convex sets ^54^. With the newly-proven mathematical theorems, we showed that the simplex of mixed expressions is a rotated and compressed version of the simplex of tissue expressions in scatter space ^55^ (Fig. 3). The vertices of the scatter simplex, characterized by the molecular markers whose expressions are maximally enriched in a particular tissue, define the optimal data-derived distinctive tissue types present in the heterogeneous samples.

CAM works by geometrically identifying the vertices (and their resident molecular markers) of the scatter simplex of globally measured expressions (Fig. 3a), *i.e.,* determining the multifaceted simplex that most tightly encloses the mixed expression profiles (Fig. 3d), and subsequently estimate the proportions and specific expression profiles of constituent subpopulations ^55^ (Fig. 3b). Tissue samples to be analyzed by CAM contain unknown number and varying proportions of molecularly distinctive (including novel) tissue types (Fig. 3e). Molecular expression in a specific tissue is modeled as being linearly proportional to the abundance of that tissue and the number of tissues present is determined by the newly-derived minimum description length (MDL) criterion ^55^.

We first eliminate proteins whose signal intensity (vector norm) is lower than 5% (noise) or higher than 95% (outlier) of the mean value over all proteins. The signals from these proteins are unreliable and could have a negative impact on the subsequent deconvolution. Second, dimension reduction is performed on the raw measurements using principal component analysis with 10 PCs. To further reduce the impact of noise/outlier data points and permit appropriate parameterization of the MDL criterion to determine the number of tissues, we aggregate protein vectors into representative clusters using affinity propagation clustering (APC) ^56-58, 22^. As an initialization-free and near-global-optimum clustering method, APC simultaneously considers all protein vectors as potential exemplars and recursively exchanges real-valued ‘messages’ between protein vectors until a high-quality set of exemplars and corresponding clusters gradually emerge. CAM detected four tissues from LAD simplex (**Fig. 3d**) and three tissues from AA simplex (**Supplemental Fig. 6c**). On the basis of the expression levels of tissue-specific marker proteins detected by CAM, the relative proportions of constituent tissues are estimated using standardized averaging that are then used to deconvolute the mixed expressions into tissue-specific profiles by non-negative least-square regression techniques^55^. Upregulated maker proteins associated with specific tissues can be detected by One-Versus-Everyone (OVO) fold change thresholding, e.g., currently set to 2 for FP tissue.

### GO Term and Pathway Enrichment Analysis

GO Term and pathway enrichment analyses using GO Term Finder ^59^, and IPA (Ingenuity Pathway Analysis, Ingenuity Systems, Redwood City, CA) tools were used to characterize the evaluable proteins obtained from the arterial samples (**Supplemental Tables 1–4**). For the GO term enrichment analysis, clustering based on semantic similarity was used to indicate which child nodes could be represented by higher level parent nodes ^60^. For the pathway analysis, the significant differentially expressed proteins with |fold change| >=1.7 and the false discovery rate (FDR) corrected p-value of 0.05 were analyzed with IPA for pathways, upstream regulators, and associated diseases and biological functions. The following parameters were used for these analyses: 1) Human genes in the Ingenuity Knowledge Base were used as the reference set; 2) Both the direct and indirect relationships were considered; 3) The confidence level was set to be “Experimentally Observed” and “High predicted” with high confidence scores; 4) The following selected tissues/cell types were used: heart, smooth muscle, fibroblasts, cardiomyocytes, endothelial cells, smooth muscle cells, and macrophages. The p-values for the identified canonical pathways, disease associations and functions were calculated using Fisher's exact test. The Benjamini-Hochberg method was used to estimate the false discovery rate, and an FDR-corrected p-value of 0.05 was used to select significantly enriched pathways. The significant upstream regulators were selected using p-value <=0.05 and |Z-score| >=2.

### Differential Dependent Network Analyses

Modeling biological networks is an important tool in systems biology to study the orchestrated activities of gene products in cells ^26^. Significant rewiring of these networks provides a unique perspective on phenotypic transitions that can occur in biological systems ^61^. Thus, instead of asking “which genes are differentially expressed”, a more interesting question is “which genes are differentially connected?” ^26,62^. To systematically characterize selectively activated or deactivated regulatory components and mechanisms, the modeling tools must effectively distinguish significant rewiring from random background fluctuations. We have developed an integrated molecular network learning method, within a well-grounded mathematical framework, to construct differential dependency networks with significant rewiring ^23, 63^. This knowledge-fused differential dependency networks (kDDN) method, implemented as a Java Cytoscape app, can be used to optimally integrate prior biological knowledge with measured data to simultaneously construct both common and differential networks ^64,65^

kDDN algorithm jointly learns the shared biological network and statistically significant rewiring across different phenotypes. Phenotype-specific data and prior knowledge are quantitatively fused via an extended Lasso model with l_1_ regularized convex optimization formulation ^64^. Based on the unique nature of the problem, we derive an efficient closed-form solution for the embedded sub-problem solved by the block-wise coordinate descent (BCD) algorithm. Since existing knowledge is often nonspecific or imperfect, kDDN uses a “minimax” strategy to maximize the benefit of prior knowledge while confining its negative impact under the worst-case scenario. Furthermore, kDDN matches the values of model parameters to the expected false positive rates on network edges at a specified significance level, and assesses edge-specific p-values on each of the differential connections ^64^.

We use kDDN to construct the network and detect significant network rewiring between FP and NL groups, where an extended Lasso-based sparse Gaussian graphic model is used to capture the network structure. The network rewiring under different phenotypes is inferred jointly by performing the BCD algorithms sequentially for all nodes. We then used permutation-based significance test to estimate p-values of the detected differential dependence edges ^63,64^. Those detected differential dependence edges with p-values larger than 0.05 were filtered out for control of false positive rate. We used different colors to distinguish differential dependency edges under each phenotype, and use solid/dashed lines to indicate positive/negative dependences between nodes. The linewidths of edges correspond to the dependency strength to give a straightforward illustration and a better comparison within the network.

## Supplemental Tables

Supplemental Table 1. Human Arterial Proteins from Left Anterior Descending (LAD) Coronary Artery (N=99) and Distal Abdominal Aortic (AA) (N=99) Samples <Data can be viewed at Peptide Atlas http://www.peptideatlas.org/PASS/PASS01066>

**Supplemental Table 2a.**
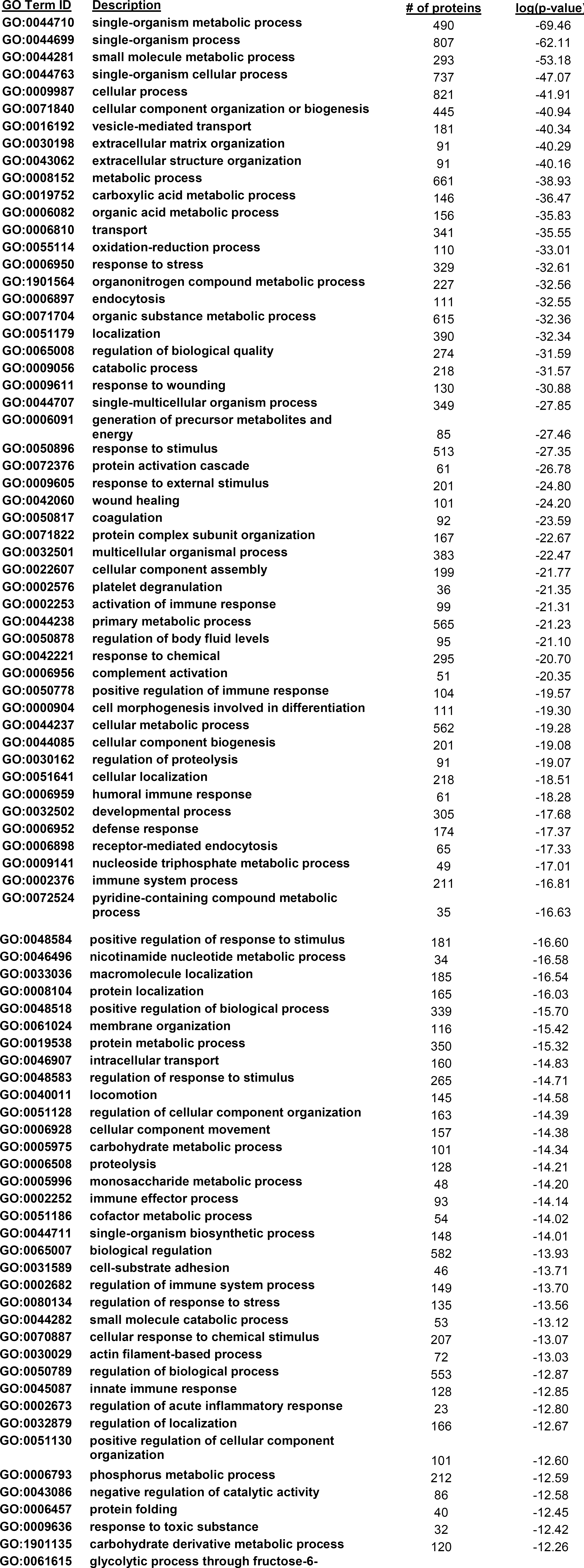
Human LAD Protein GO Terms: Biologic Processes (top 100 GO Terms)

**Supplemental Table 2b.**
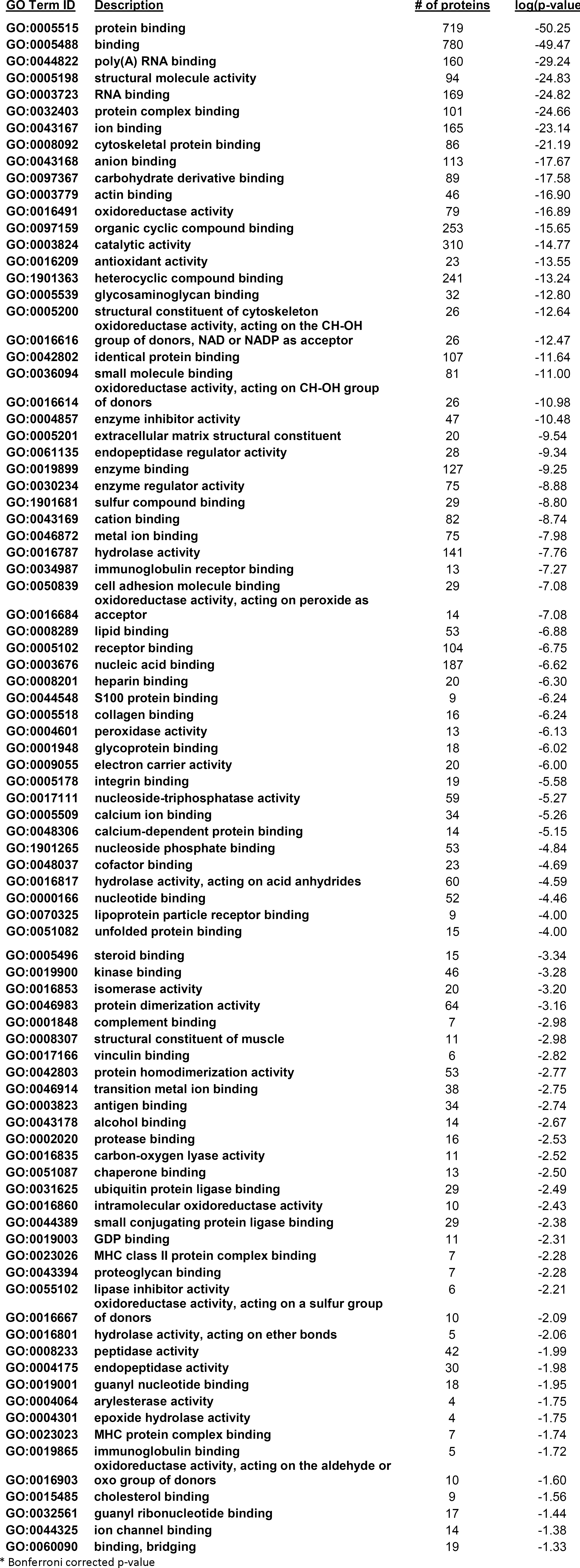
Human LAD Protein GO Terms: Molecular Functions (adj. p-value < 0.05*)

**Supplemental Table 2c.**

Human LAD Protein GO Terms: Cellular Component (adj. p-value < 0.05*)

**Supplemental Table 3a.**
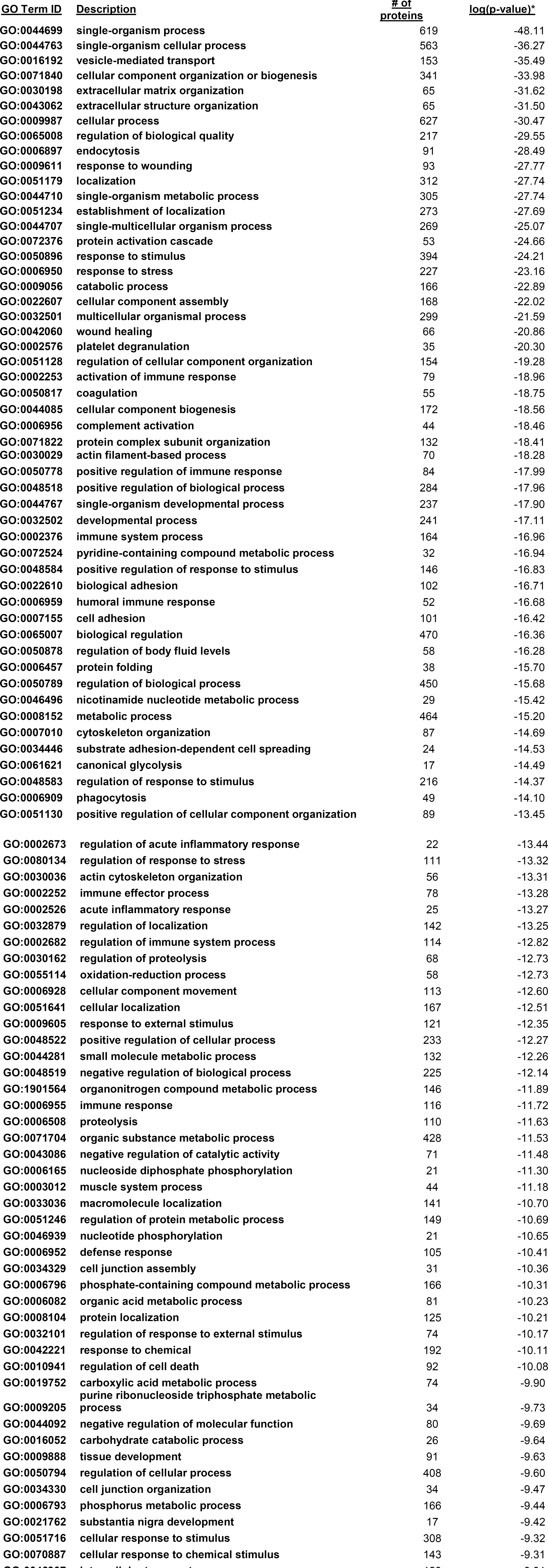
Human Distal Aortic Protein GO Terms: Biologic Process (top 100 GO Terms)

**Supplemental Table 3b.**
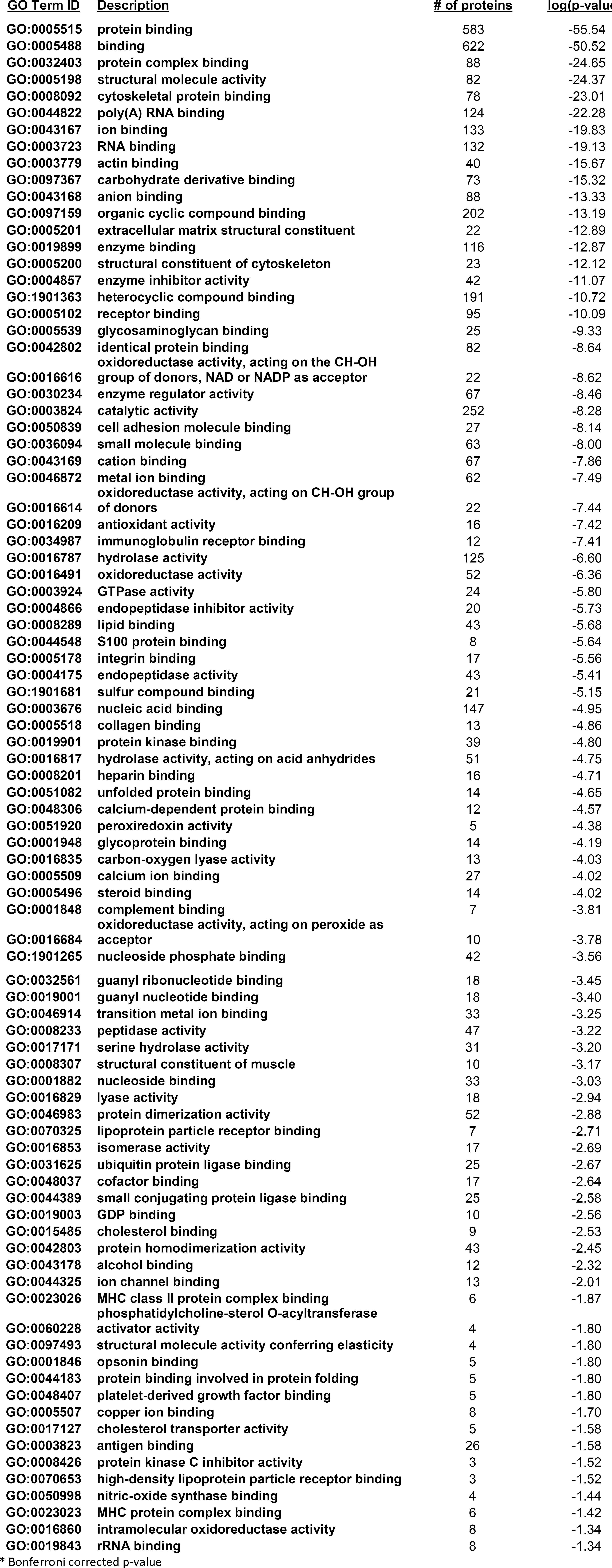
Human Distal Aortic Protein GO Terms: Molecular Function (adj. p-value < 0.05*)

**Supplemental Table 3c.**
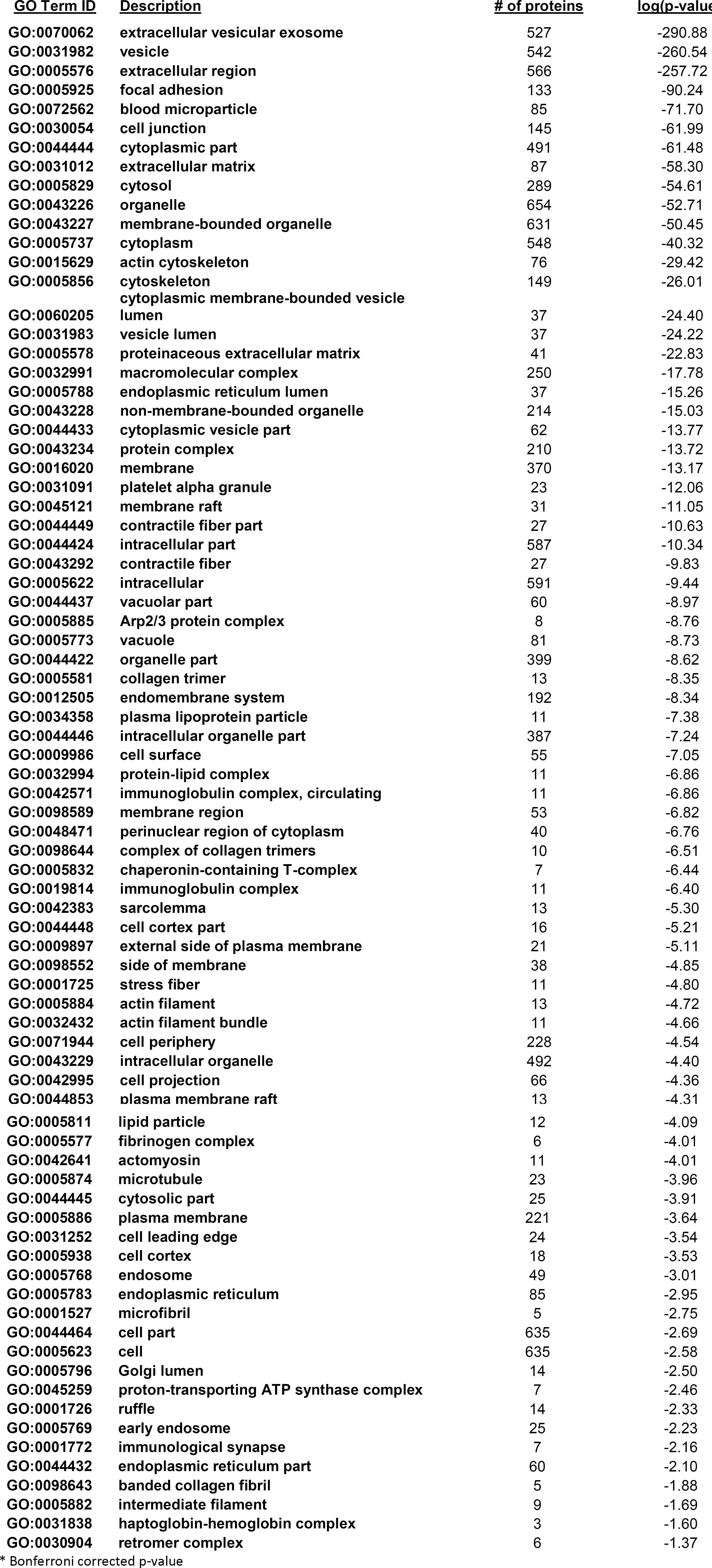
Human Distal Aortic Protein GO Terms: Cellular Component (adj. p-value < 0.05*)

**Supplemental Table 4.**
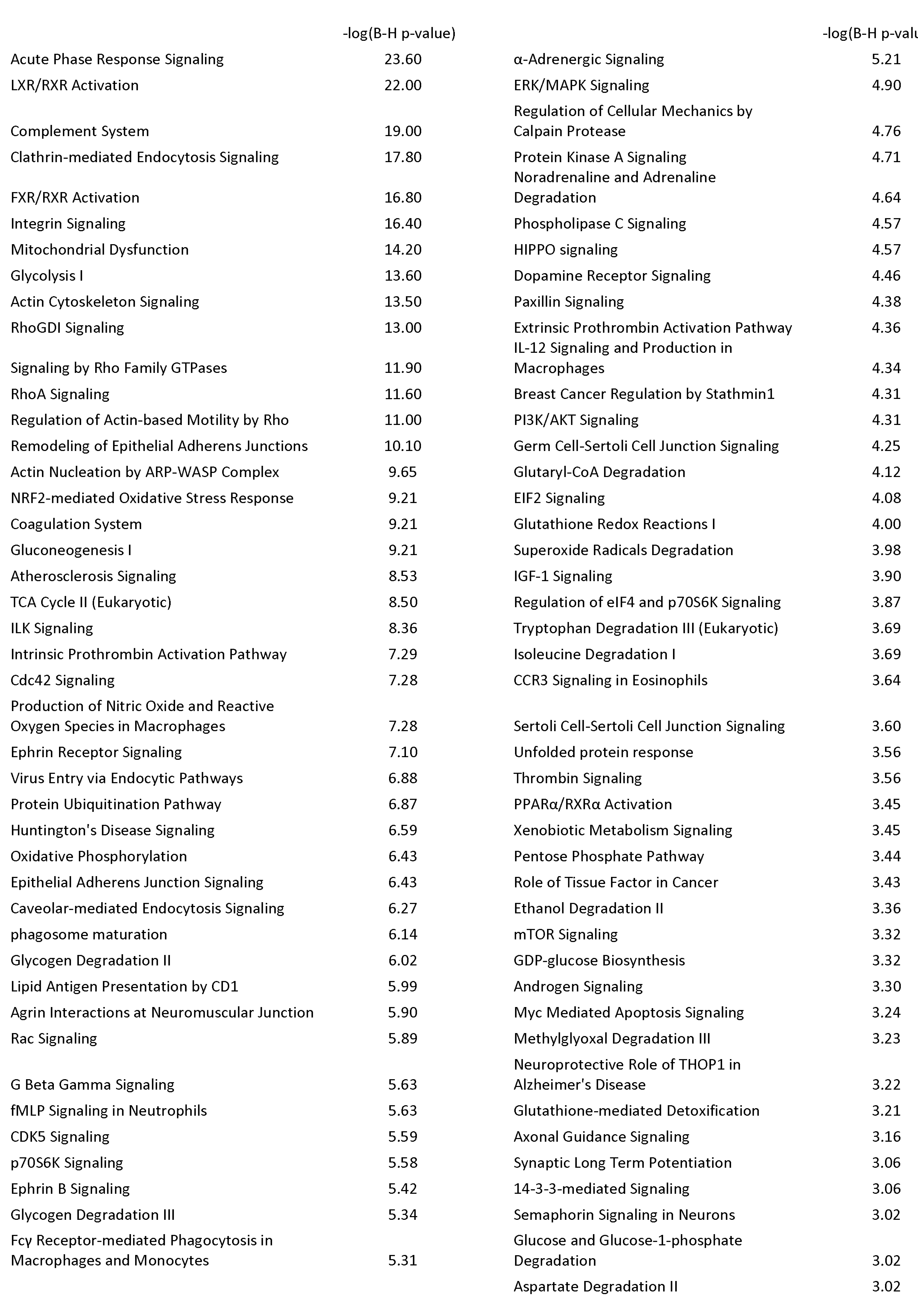
Enriched Ingenuity Canonical Pathways Among 944 Human LAD Proteins (FDR<0.001)

**Supplemental Table 5.**
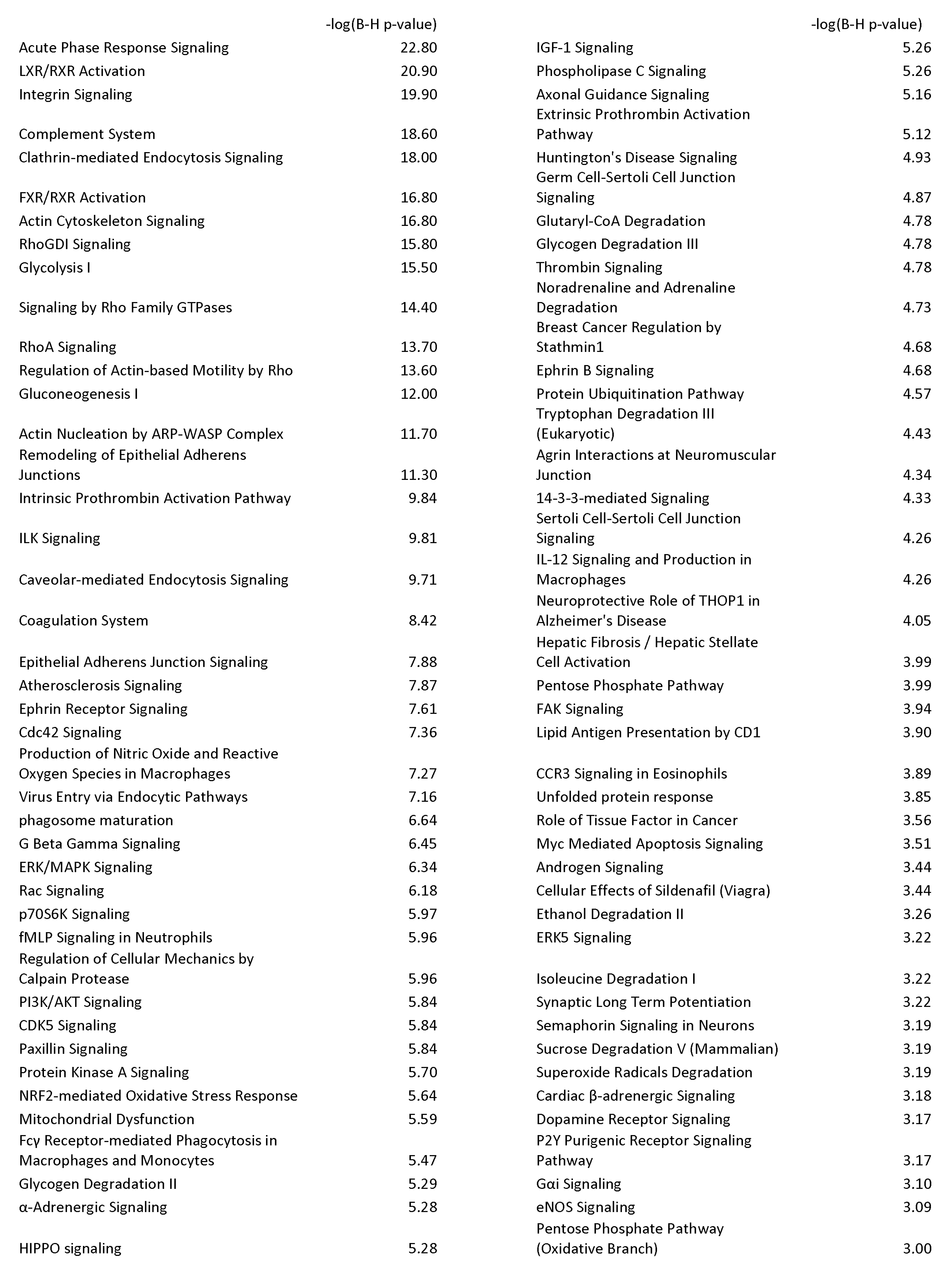
Enriched Ingenuity Canonical Pathways among 725 Human Distal Aortic Proteins (FDR<0.001)

**Supplemental Table 6.**
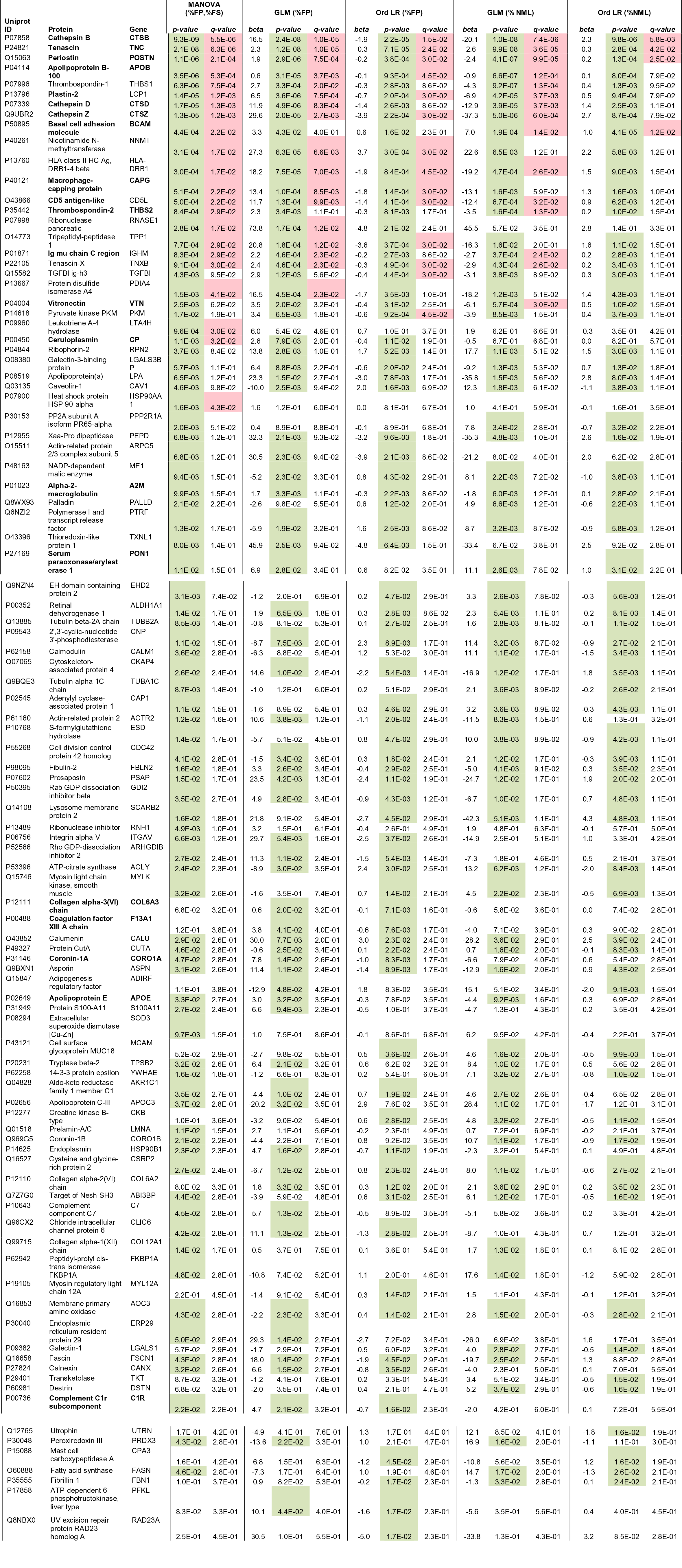
Regression Models of Association Between Individual Proteins and Extent of Surface Area Involved* with Fibrous Plaque (FP) or Normal Intima (NL) in Human LAD Arterial Samples (top 100 proteins). In models of %FP, the r FP% fraction includes fatty streaks (FS) and NML intima. Likewise, in models of % NL, the non-NL faction includes both FP ar FS. Accordingly, we modeled both %FP and %NL separately. We also classified samples into three level ordinal categories of extent of FP and extent of NL and performed ordinal logistic regression (Ord LR). This approach is less statistically powerful, makes fewer distributional assumptions about the measures of extent of disease. In addition, we used MANOVA to jointly nr the distribution of %FP, %FS and %NL. P-values and q-values ≤0.05 are highlighted in green and pink respectively. **Proteins i bold (n=21) are also included in the top 100 AA proteins**

**Supplemental Table 7.**
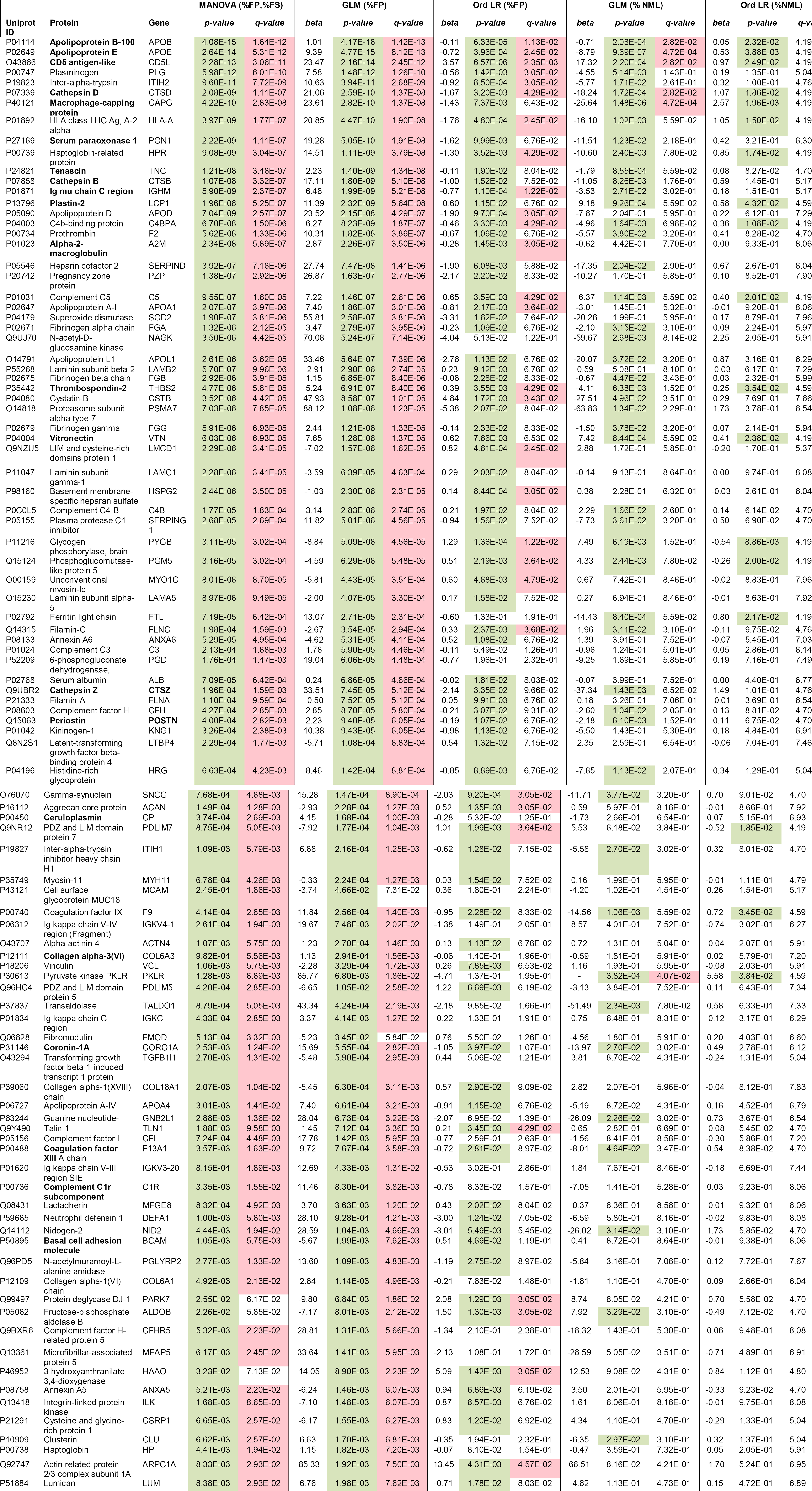
Regression Models of Association Between Individual Proteins and Extent of Surface Area Involvei with Fibrous Plaque (FP) or Normal (NML) in Human Abdominal Aorta Samples (top 100 proteins). In models of %FP, the r FP% fraction includes fatty streaks (FS) and NML intima. Likewise, in models of % NML, the non-NML faction includes both F and FS. Accordingly, we modeled both %FP and %NML separately. We also classified samples into three level ordinal categi of extent of FP and extent of NML and performed ordinal logistic regression (Ord LR). This approach is less statistically powf but makes fewer distributional assumptions about the measures of extent of disease. In addition, we used MANOVA to joint model the distribution of %FP, %FS, and %NML. P-values and q-values ≤0.05 are highlighted in green and pink respectively. **Proteins in bold (n=21) are also included in the top 100 LAD proteins.**

**Supplemental Table 8.**
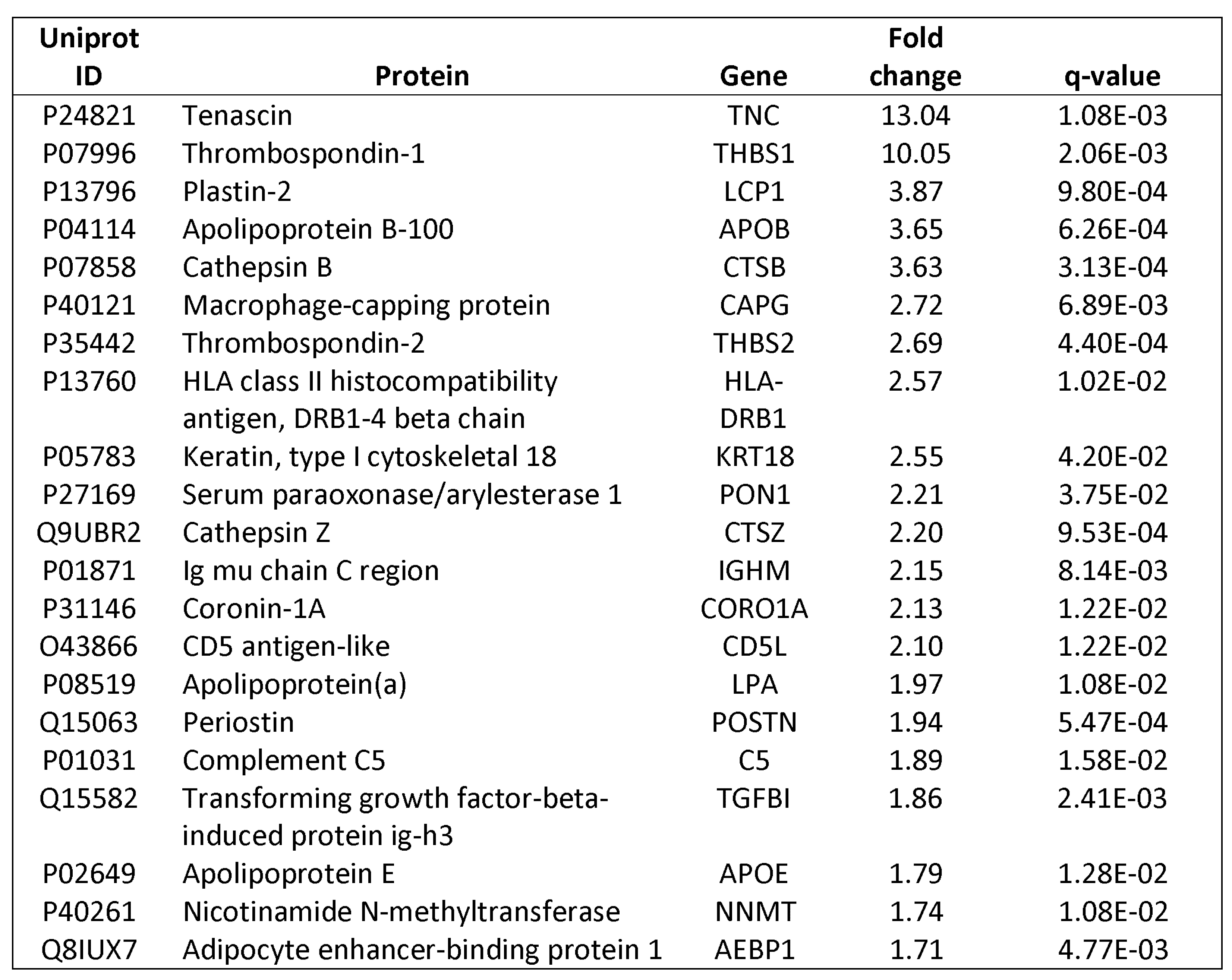
Up-Regulated Proteins in LAD Fibrous Plaque Samples (n=15) Compared with Normal LAD Samples (n=30). Samples were selected using patho-proteomic phenotyping designed to integrate pathologist assessment of extent of disease and CAM deconvolution of global protein profiles (fold-change = mean FP/ mean NL).

**Supplemental Table 9.**
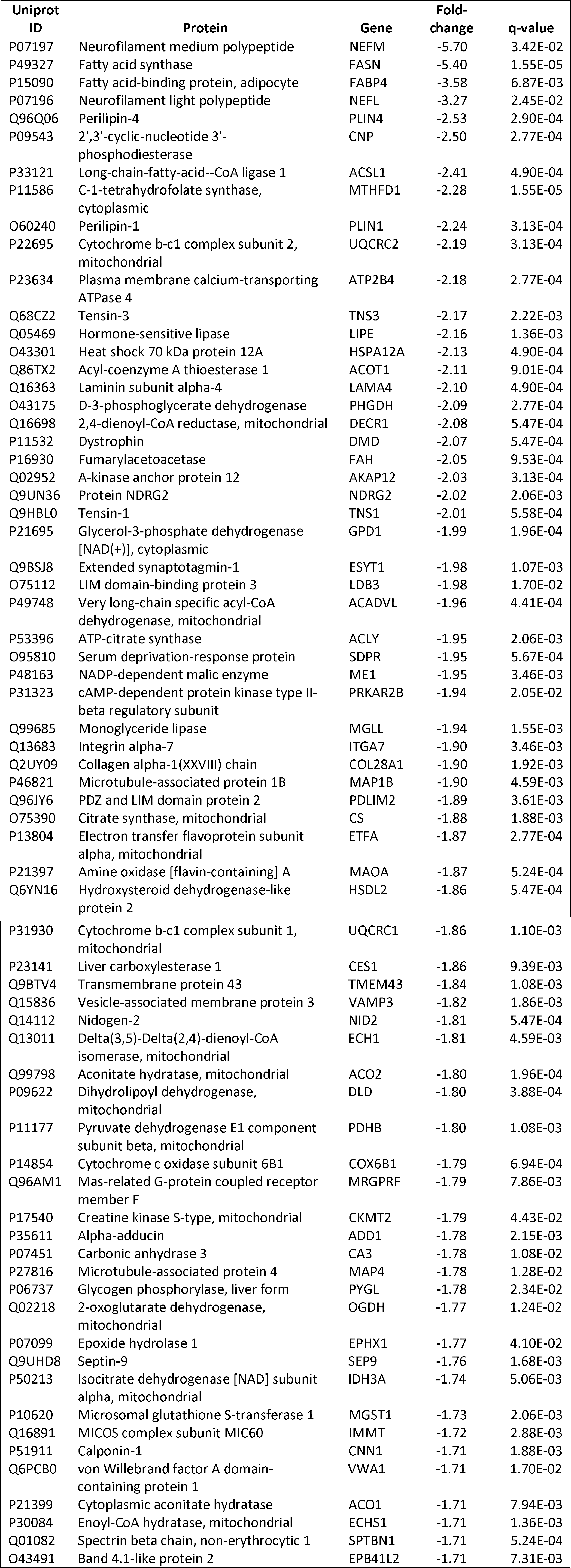
Down-Regulated Proteins in LAD Fibrous Plaque Samples (n=15) Compared with Normal LAD Samples (n=30). Samples were selected using patho-proteomic phenotyping designed to integrate pathologist assessment of extent of disease and CAM deconvolution of global protein profiles. (For down regulated proteins Fold-change =-1* mean NL/mean FP.)

**Supplemental Table 10.**
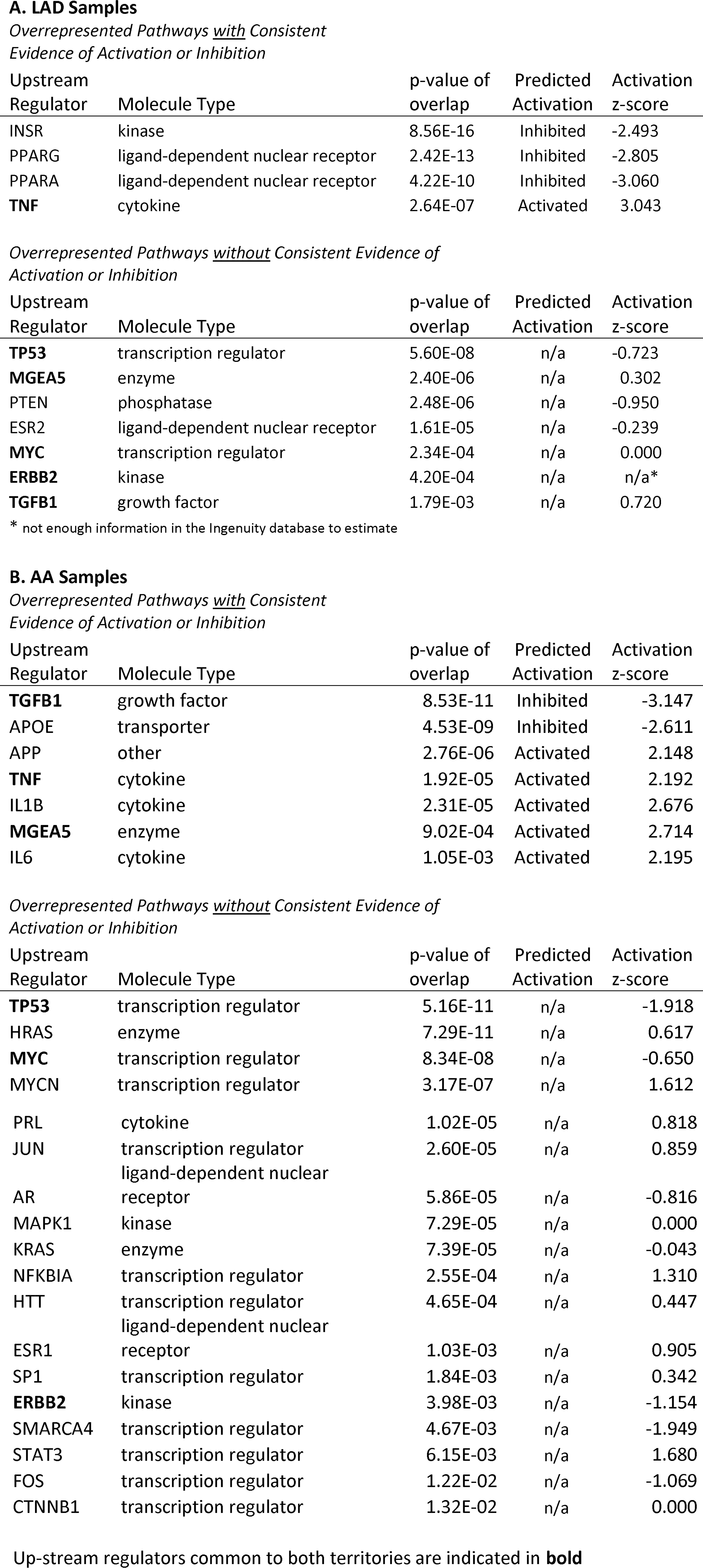
Predicted Upstream Master Regulators for Up-and Down-Regulated Fibrous Plaque Proteins A. LAD Samples. Samples were selected using patho-proteomic phenotyping designed to integrate pathologist assessment of extent of disease and CAM deconvolution of global protein profiles. (For down regulated proteins Fold-change + −1% mean NL/mean FP.)

**Supplemental Table 11.**
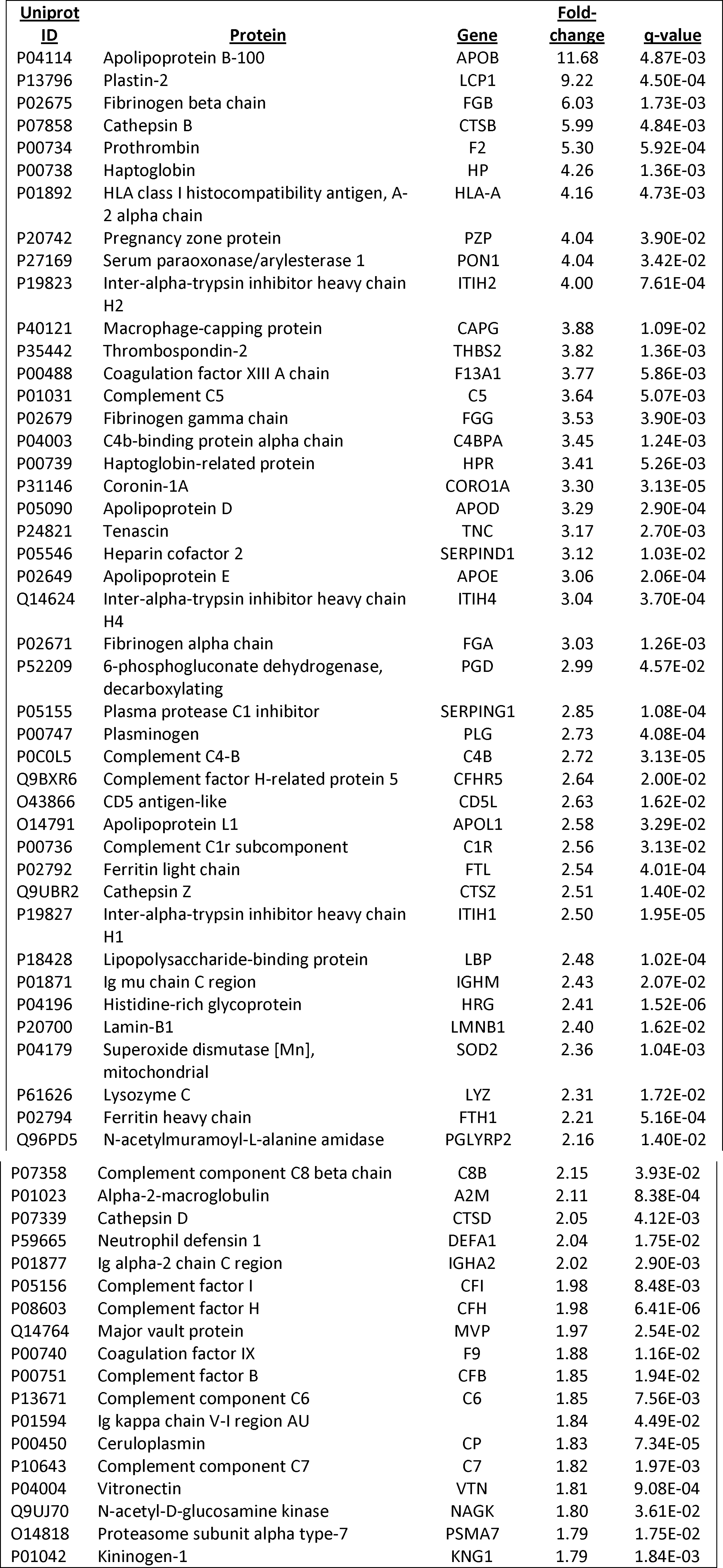
Up-Regulated Proteins in Fibrous Plaque Enriched AA Samples (n=7) Compared with Normal AA Samples (n=15). Samples were selected using patho-proteomic phenotyping designed to integrate pathologist assessment of extent of disease and CAM deconvolution of global protein profiles (fold-change = mean FP/ mean NL).

**Supplemental Table 12.**
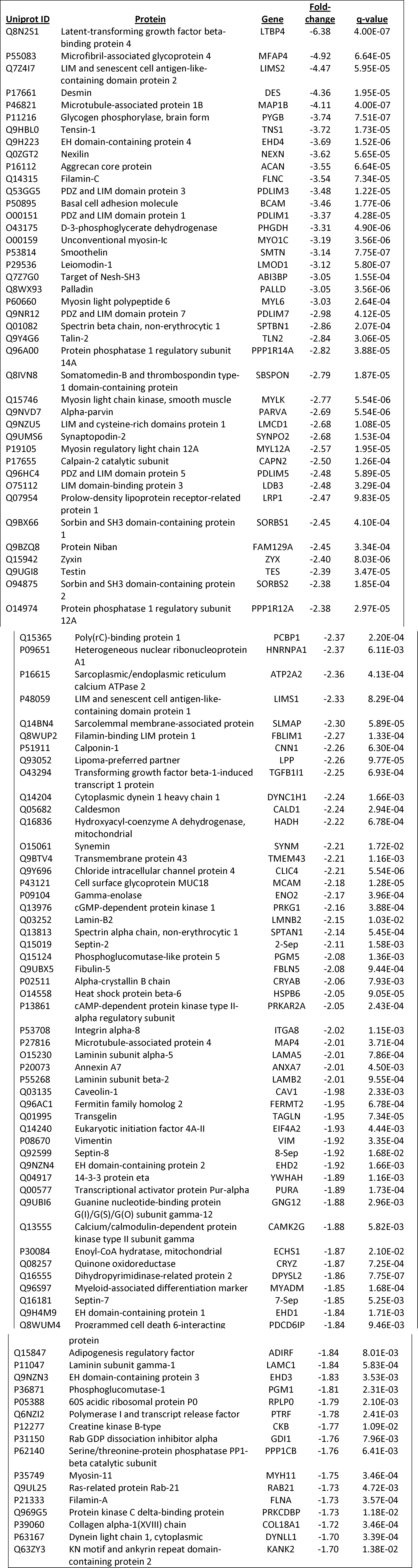
Down-Regulated Proteins in Fibrous Plaque Enriched AA Samples (n=7) Compared with Normal AA Samples (n=15). Samples were selected using patho-proteomic phenotyping designed to integrate pathologist assessment of extent of disease and CAM deconvolution of global protein profiles. (For down regulated proteins Fold-change =-1* mean NL/mean FP)

**Supplemental Table 13.**
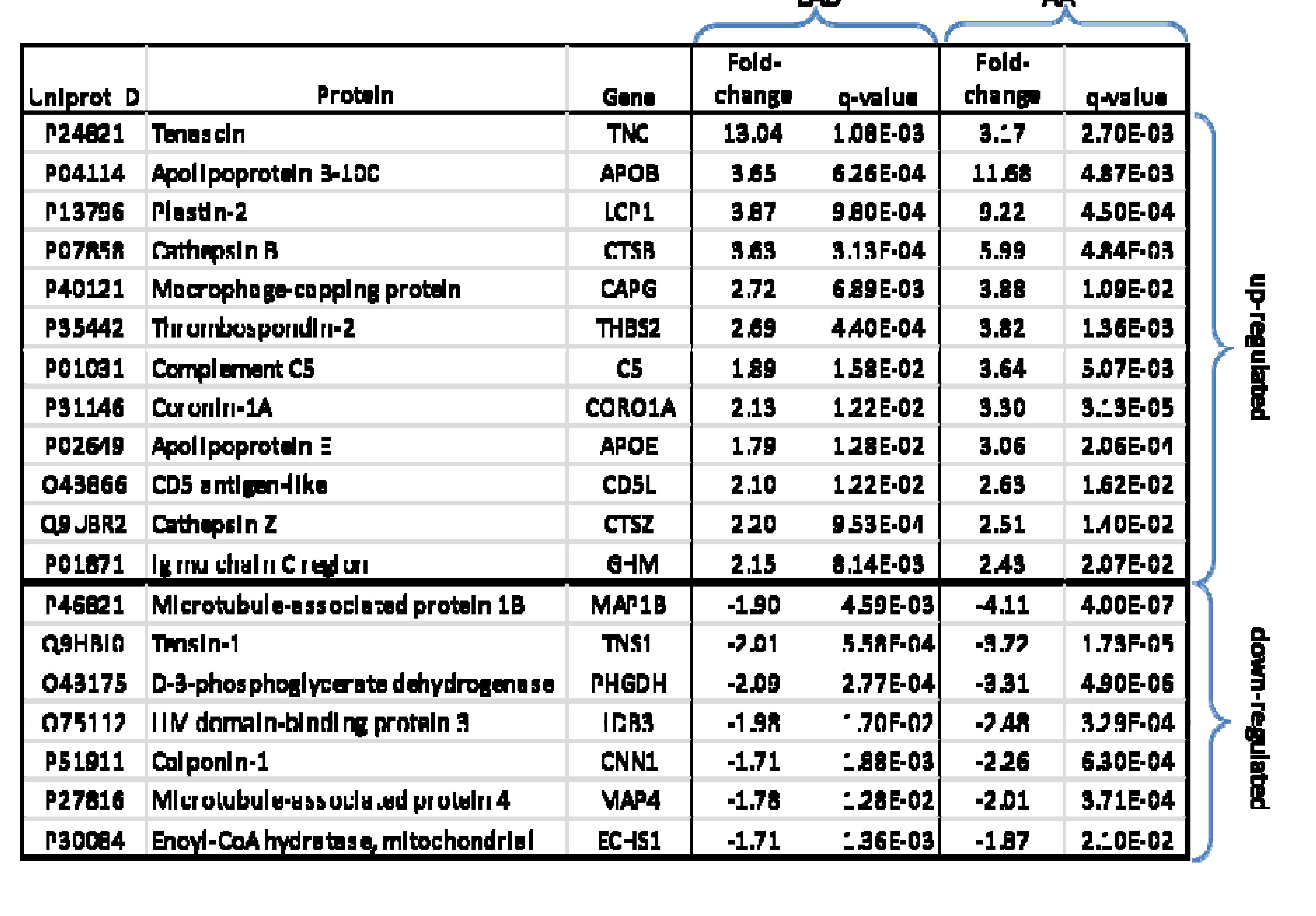

## Supplemental Figures

**Supplemental Figure 1a.**
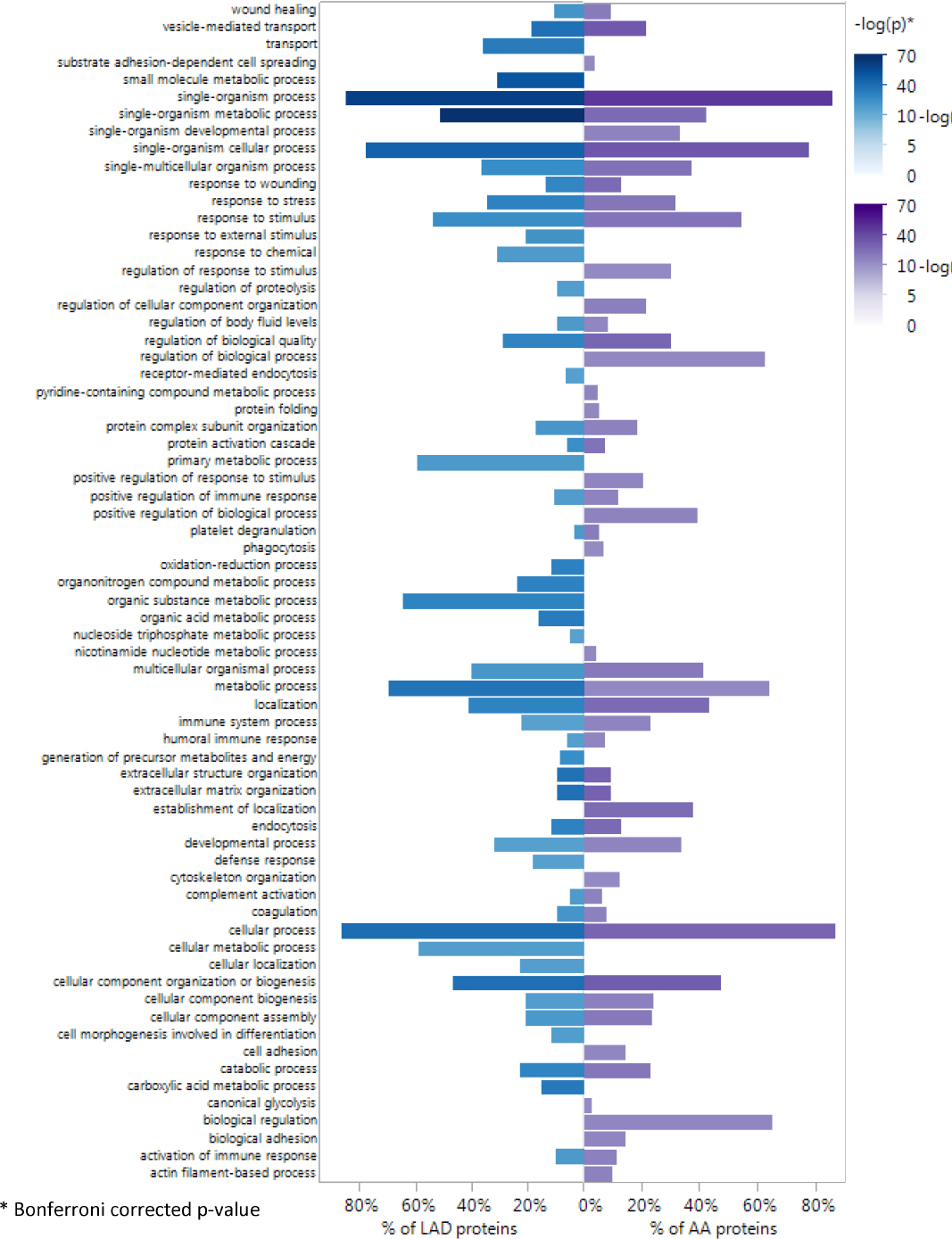
Enriched GO:Biological Process Terms From Human LAD and AA Proteins. GO enrichment analysis was performed using 944 LAD proteins (left half of the plot-blue) and 725 AA proteins (right half of the plot-purple) that were included for subsequent analysis. GO terms with Bonferroni adjusted p-value <0.05 in either anatomic territory are presented.

**Supplemental Figure lb.**
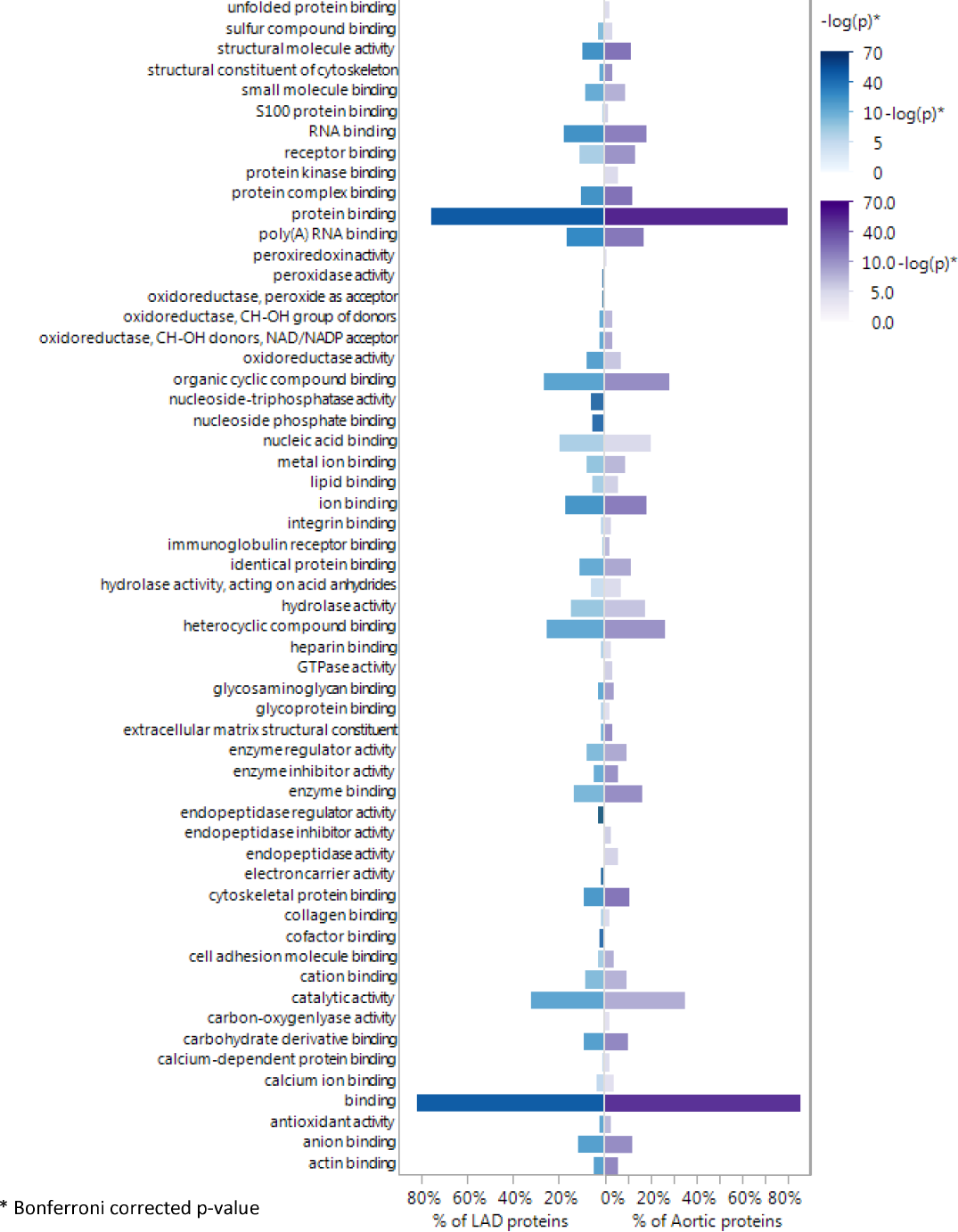
Enriched GO:Molecular Function Terms From Human LAD and AA Proteins. GO term enrichment analysis was performed using 944 LAD proteins (left half of the plot-blue) and 725 AA proteins (right half of the plot-purple) that were included for subsequent analysis. GO terms with Bonferroni adjusted p-value <0.05 in either anatomic territory are presented.

**Supplemental Figure lc.**
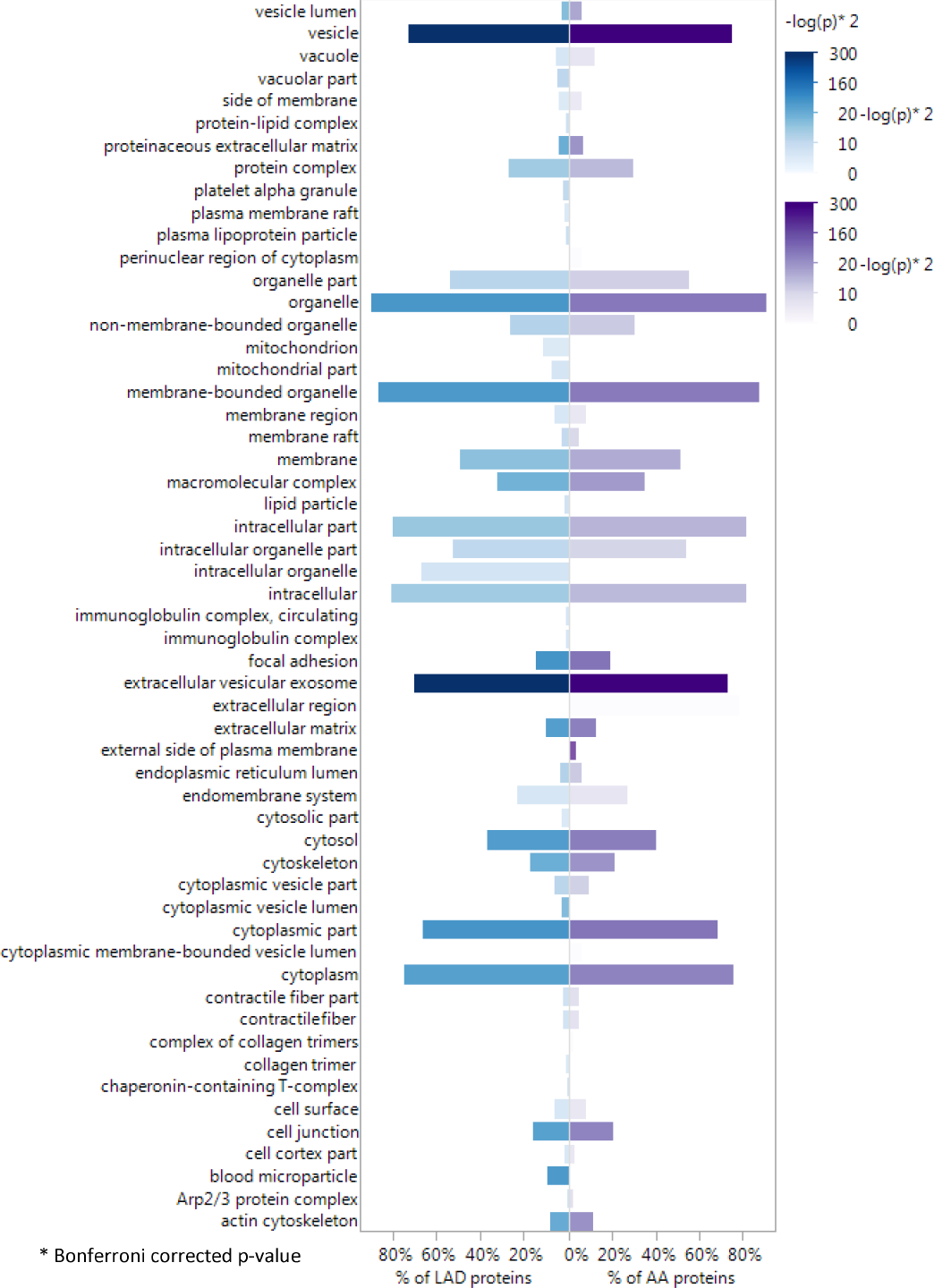
Enriched GO:Cellular Component Terms From Human LAD and AA Proteins. GO term enrichment analysis was performed using 944 LAD proteins (left half of the plot-blue) and 725 AA proteins (right half of the plot-purple) that were included for subsequent analysis. GO terms with Bonferroni adjusted p-value <0.05 in either anatomic territory are presented.

**Supplemental Figure 2.**
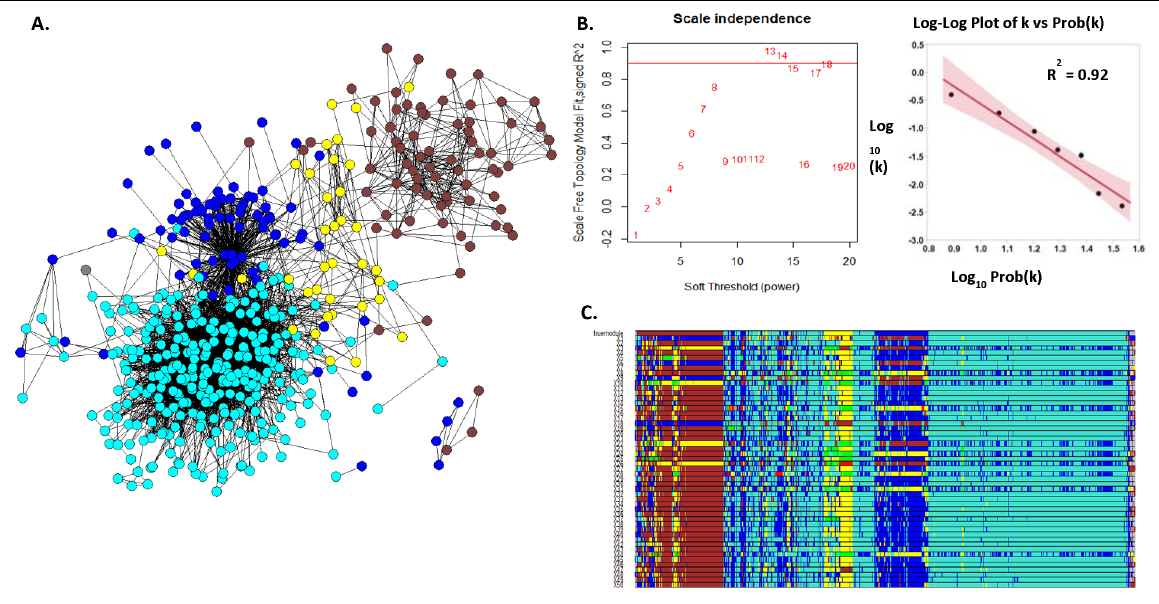
Weighted Co-Expression Network Analysis of Human Abdominal Aortic (AA) Proteins. A. Adjacency map of AA proteins color coded by module assignment based on hierarchical clustering of the topological overlap matrix (TOM)-based dissimilarity measure. For clarity of presentation only nodes (proteins) with at least one edge (adjacency measure (k)) > 98%tile are shown. B. Left panel shows that scale-free topology is well approximated when the adjacency power parameter (3= 8. Right panel shows the log-log plot of adjacency (k) vs prob(k) with (3= 8, supporting a power-law relationship in the connectivity of the expressed proteins. C. Module assignment for fifty 90% random samples of the data illustrating the overall stability of the modular structure of the protein expression patterns. Colors are assigned according cluster size which may vary with each random sample. As a result actual color assignment may vary from run to run, but module membership remains relatively stable.

**Supplemental Figure 3.**
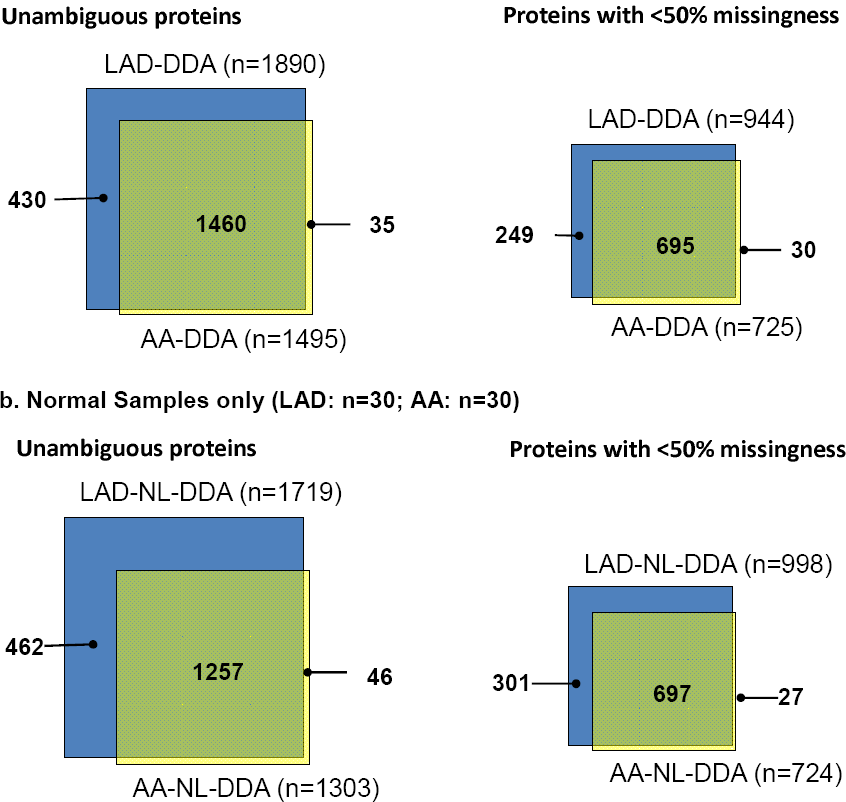
Number of identifiable proteins from human LAD and AA arterial
samples. a. All samples (n=99 samples from each territory). Left Venn diagram: proteins detected at least once in any sample according to anatomic territory; Right Venn diagram: proteins detected in >50% of the samples. GO term analysis of the LAD only proteins indicated significant enrichment of mitochondrial proteins (p-values: unambiguous proteins = 1.8e-28; proteins with <50% missingness = 1.7e-6) b. Nomal samples only (n=30 from each territory). Left Venn diagram: proteins detected at least once in any sample according to anatomic territory; Right Venn diagram: proteins detected in >05% of the samples. GO term analysis of the LAD only proteins indicated significant enrichment of mitochondrial proteins (p-values: unambiguous proteins = 1.0e-13; proteins with <50% missingness p-value = 1.9e-12)

**Supplemental Figure 4.**
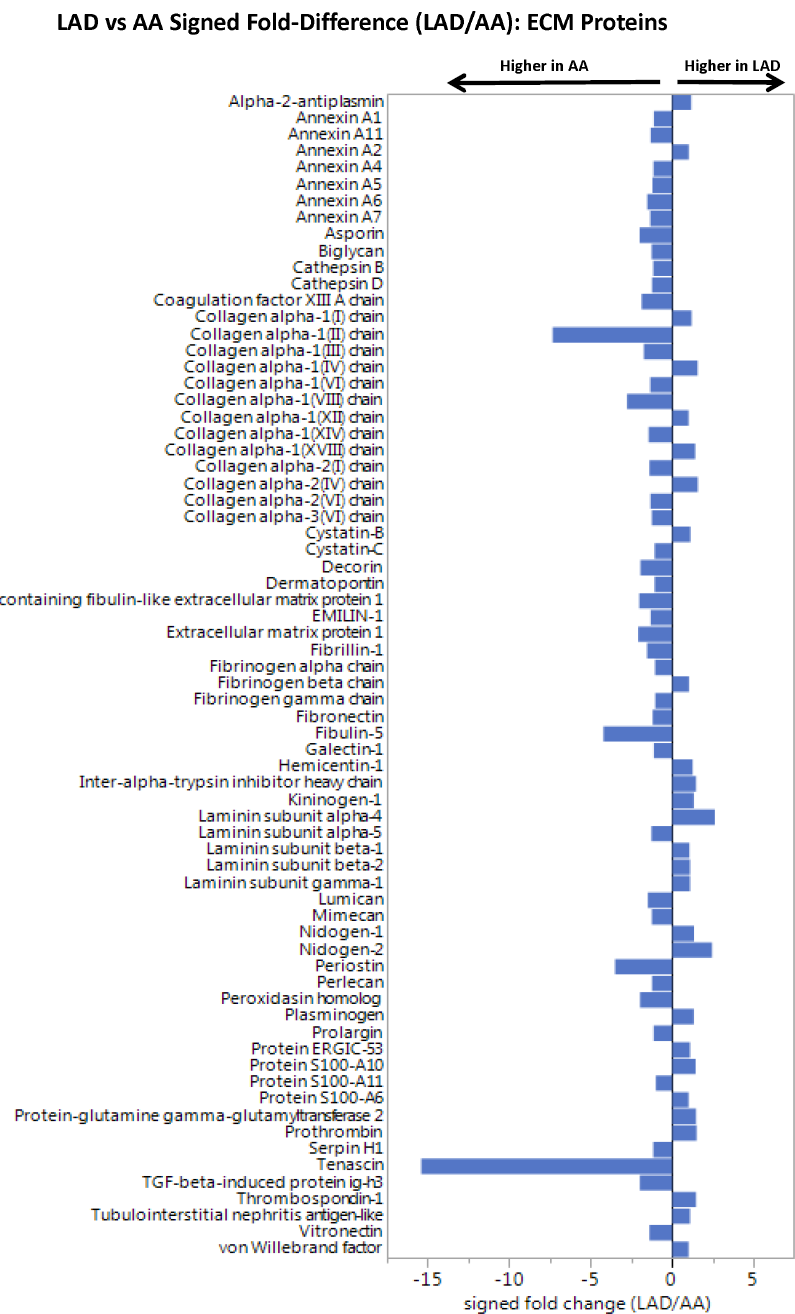
Comparison of Mitochondrial Proteins in Normal LAD and AA Samples. Data independent MS (SWATH) analysis of completely normal LAD and AA samples (n=15 from each anatomic location) with adjustment for age, sex, MYH11, RABA7A, TERA, G6PI. Mean fold-difference = 6% (AA>LAD), p= 0.01.

**Supplemental Figure 5.**
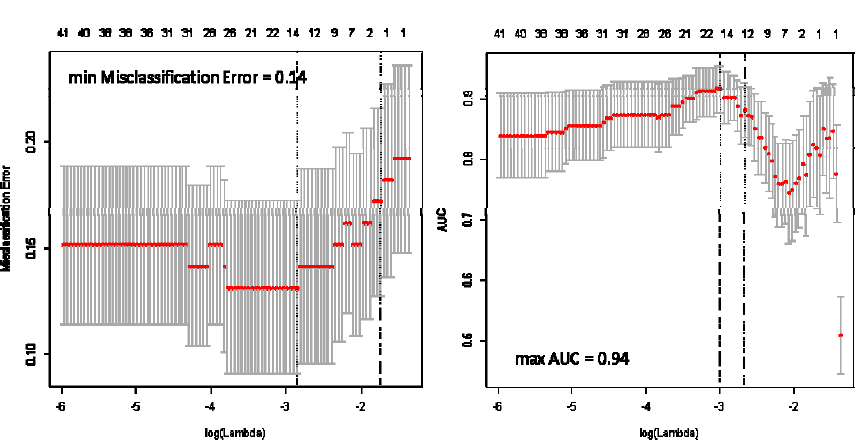
Elastic Net Modelling of Proteins Predicting Presence or Absence of Fibrous Plaque in LAD (N=99) Samples. Graphs: Misclassification and AUC across a range of lambdas for two typical elastic net models predicting presence of fibrous plaque. Red points indicate the mean and bars indicate the SE for the estimate of misclassification or AUC across 10-folds in a cross-fold validation scheme. Numbers across the top indicate the number of variables (proteins) remaining in the model as the model progresses fi the most inclusive (left) to the most parsimonious (right).

**Table:**
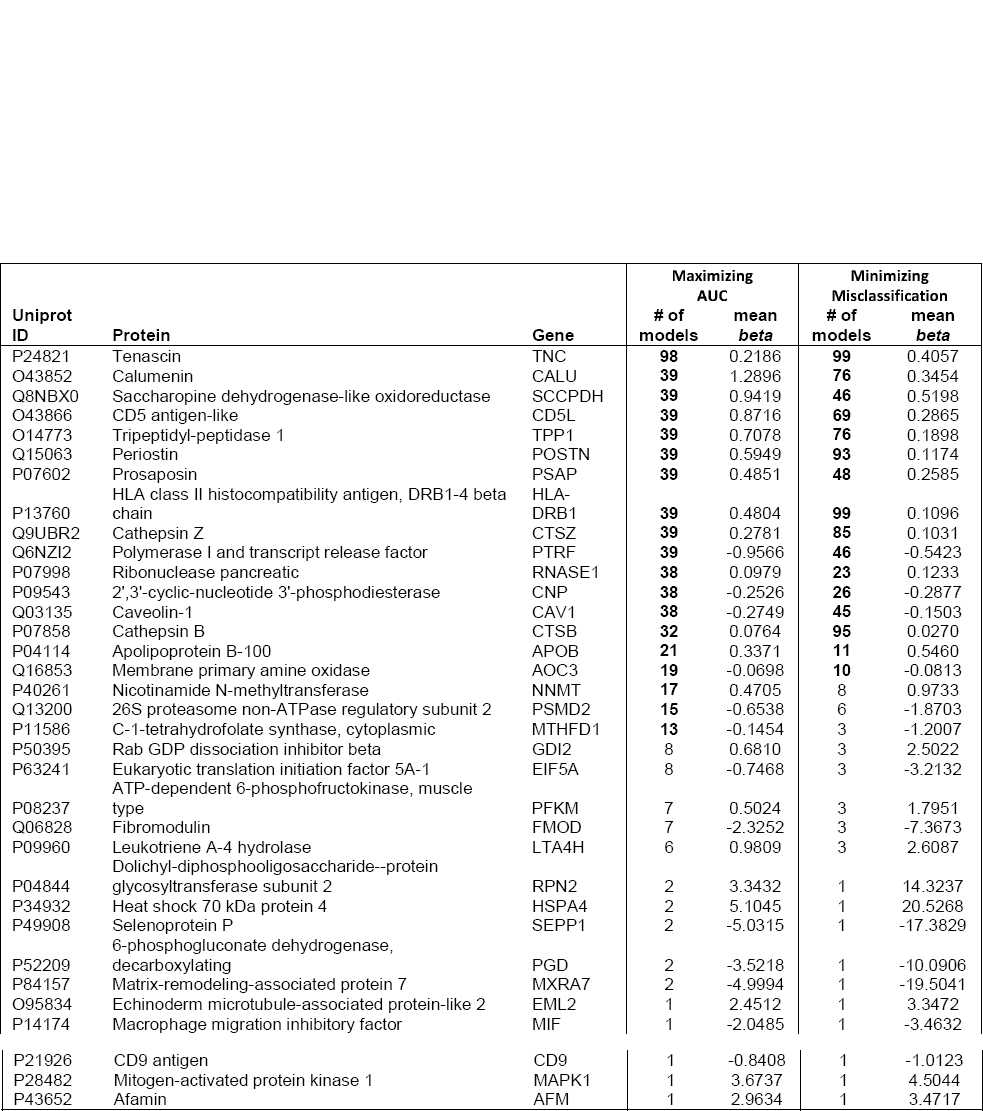
Models were developed using crossfold validation to select the lambdas optimizing AUC or misclassification. To minimize the effect of chance in selecting folds for the cross-fold validation each model was run 100 times and the selected proteins and their respective beta coefficients were recorded. The table indicates the number of times each protein was selected in each of 100 models and the mean beta for the subset of models where the protein in question was included. Proteins included in 10 or more models are indicated by bold in the # of models columns. Several highly correlated proteins competed for entry into these models.

**Supplemental Figure 6.**
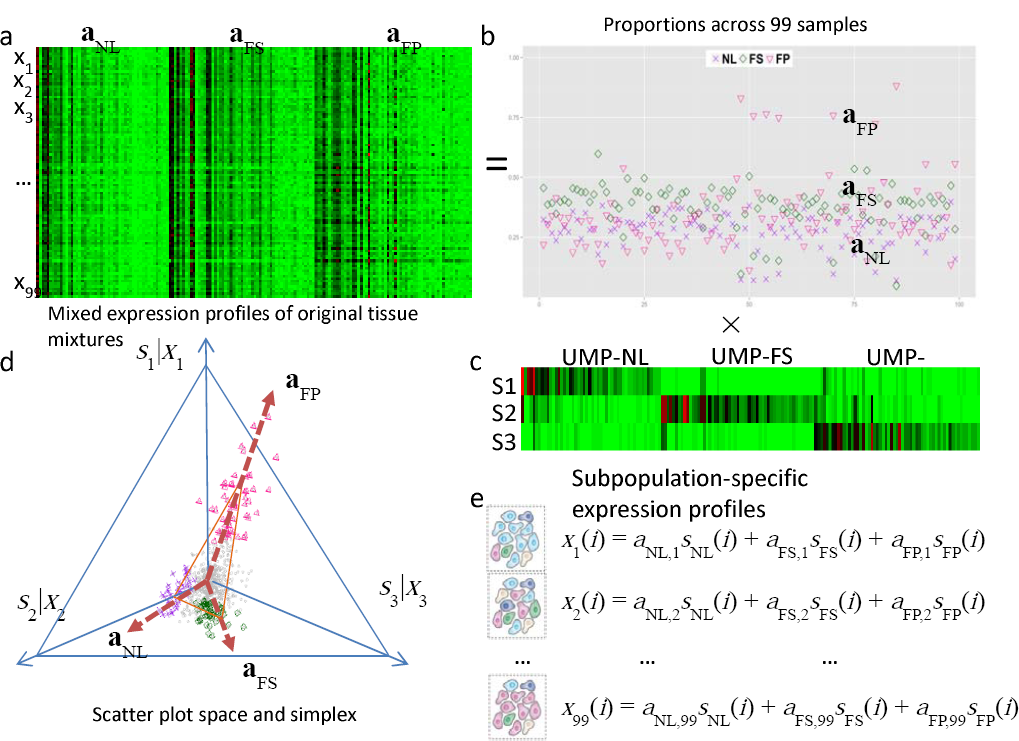
Convex Analysis of Mixtures of AA Protein Data. a. Heatmap of mixed expressions of upregulated marker proteins (UMP) in 99 AA samples, b. Estimated proportions of NL, FS, and FP across 99 AA samples. Note: The AA data only supported identification of three vertices compared with four vertices in the LAD data. c. Heatmap of subpopulation-specific expressions of upregulated marker proteins, d. Geometry of the mixing operation in scatter space that produces a compressed and rotated scatter simplex whose vertices host subpopulation-specific upregulated marker proteins and correspond to mixing proportions, e. Mathematical description on the protein expression readout of multiple distinct subpopulations.

**Supplemental Figure 7.**
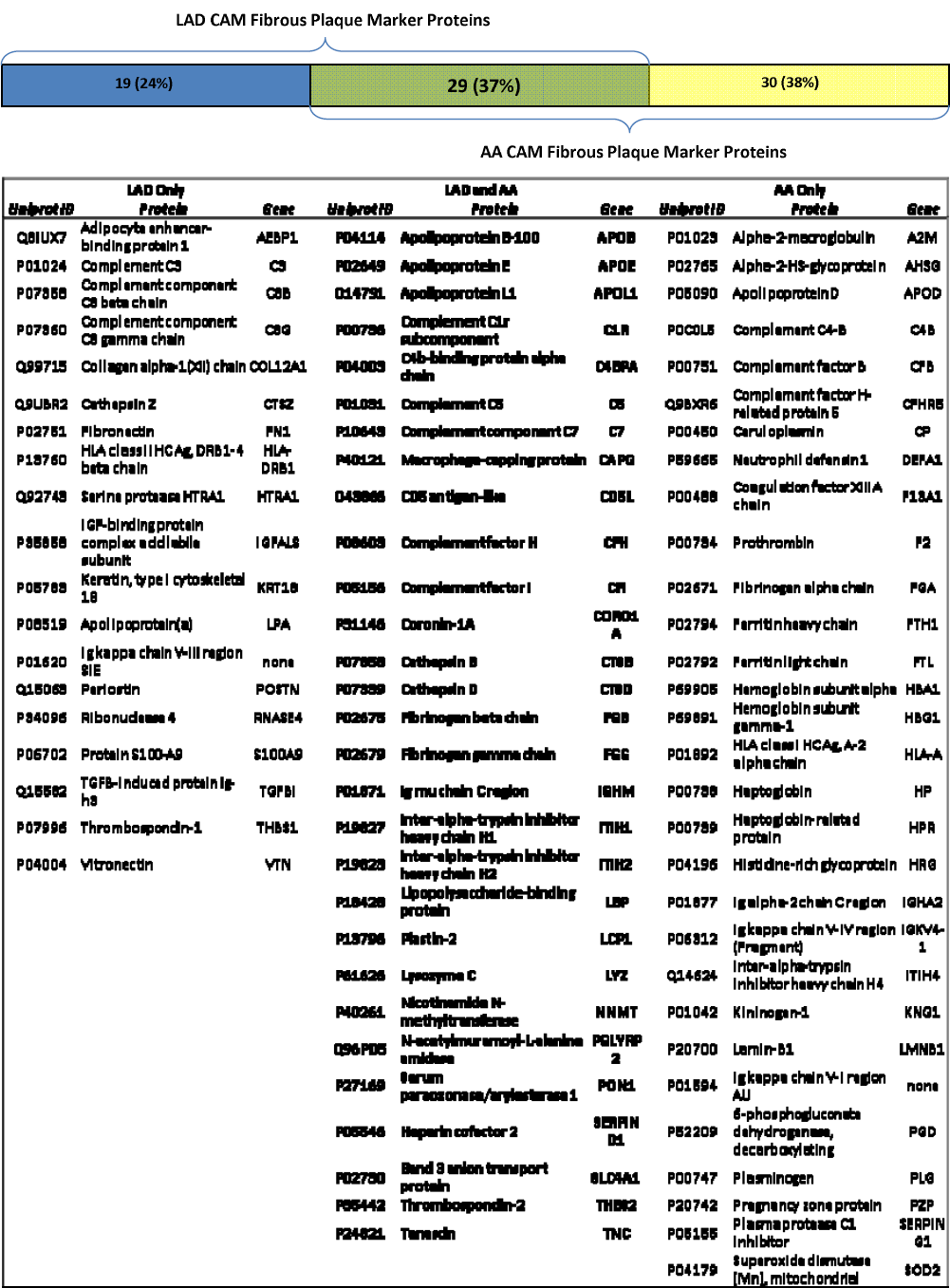
Convex Analysis of Mixtures (CAM) Identified Upregulated Fibrous Plaque Marker Proteins for LAD (n=99) and AA (n=99) Samples.

**Supplemental Figure 8.**
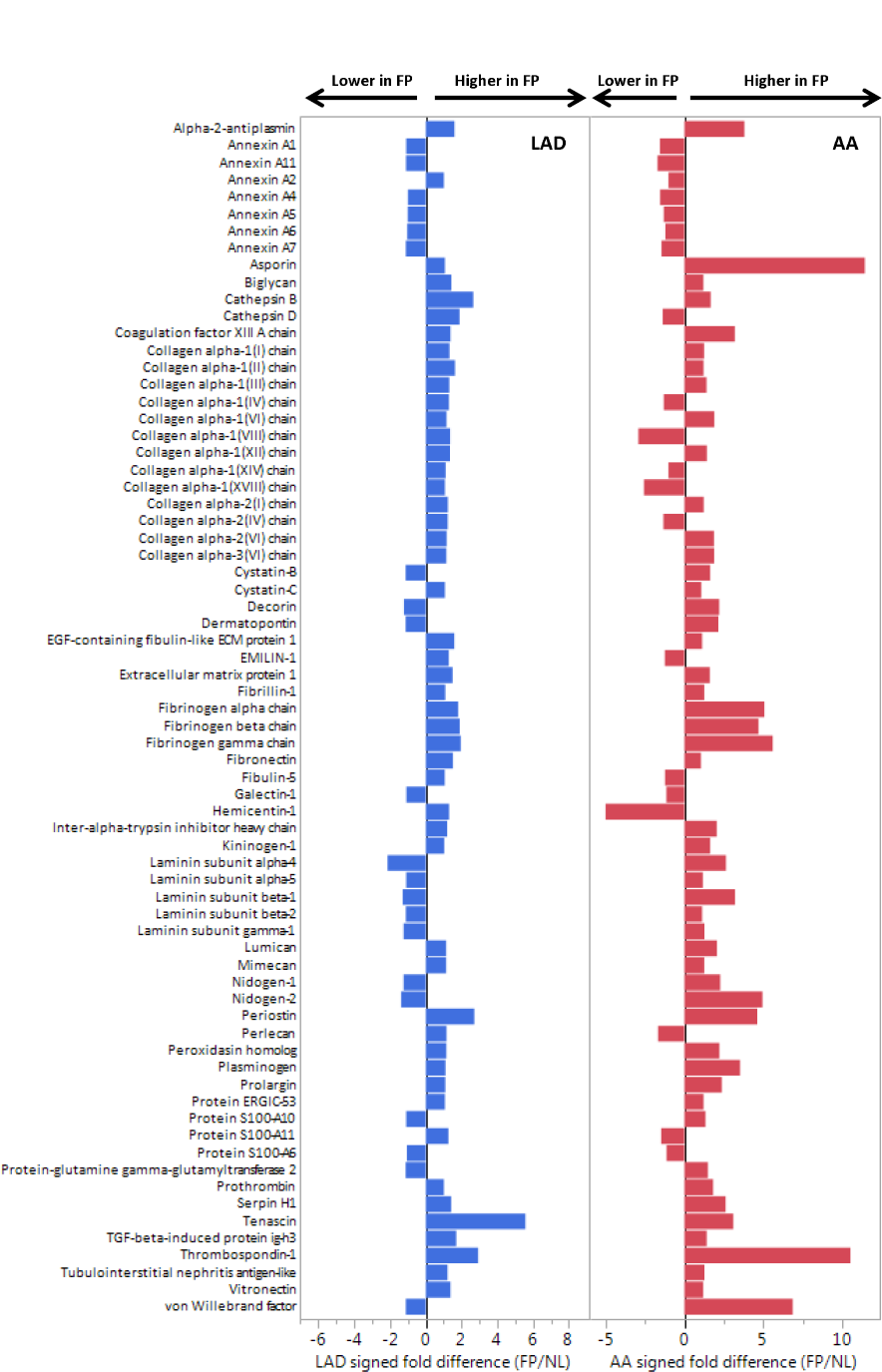
Comparison of Extracellular Matrix Proteins in FP vs NL Samples in LAD and AA Samples. Data independent acquisition (SWATH) analysis was used to compare a targeted set of mitochondrial proteins in LAD (FP n=15; NL n=30) and AA (FP n=9; NL n=18) samples after adjustment for age, sex, MYH11, RABA7A, TERA, G6PI. Histogram bars indicate relative difference between FP and NL samples in each anatomic location. LAD Mean FP/NL = 1.25, MANOVA p-value = 0.02; AA Mean FP/NL = 1.78, p-value = 0.017

**Supplemental Figure 9.**
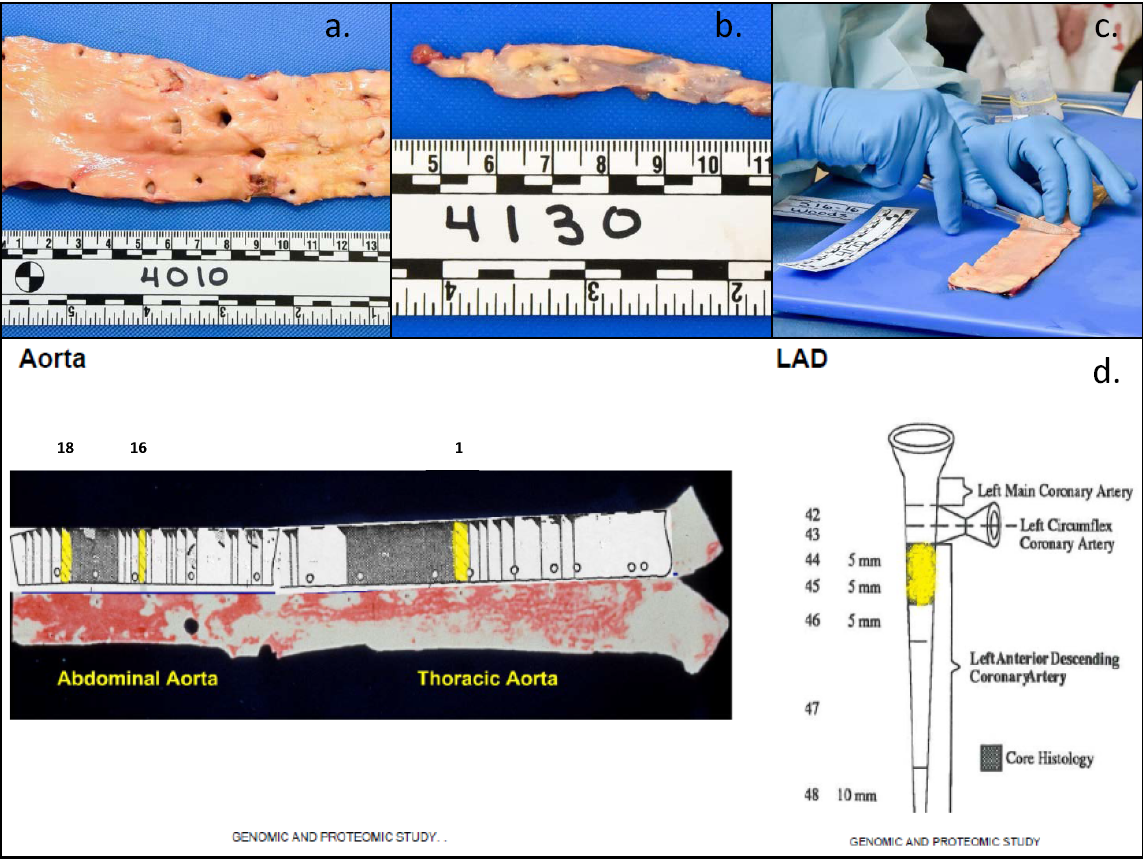
Arterial Sample Acquisition: **a**.) distal aorta; **b.**) left anterior descending (LAD) coronary artery; **c.**) collection of a gram of tissue from the distal aortic specimen; **d.**) Standardized locations for sample collection. Specimens were obtained from Sections 1,16,18, 44 and 45 (indicated in yellow) for all autopsies. These locations correspond to locations originally established and used for the Pathobiological Determinants of Atherosclerosis in Youth study ^29^.

**Supplemental Figure 10.**
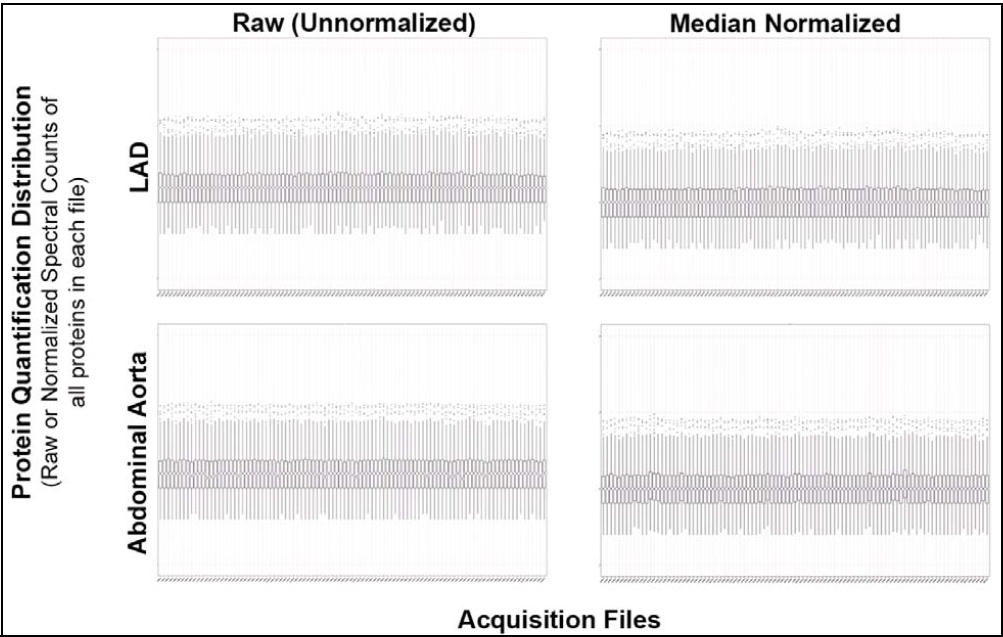
Acquisition and Protein Quantification Quality Control (QC). QC was assessed by plotting the distribution in the form of a box and whisker plot of protein quantitation values for all identified proteins across all acquired specimen. Notches represent the 95% confidence interval of the median, and files with unaligned notches represent substantially different quantitative distributions. Using this approach, while the LAD files all had nicely aligned quantitative distributions of raw spectral counts with overlapping medians for all files (upper left panel), the aorta samples had some acquisitions with uneven quantitative distribution. For aorta

**Supplemental Figure 11.**
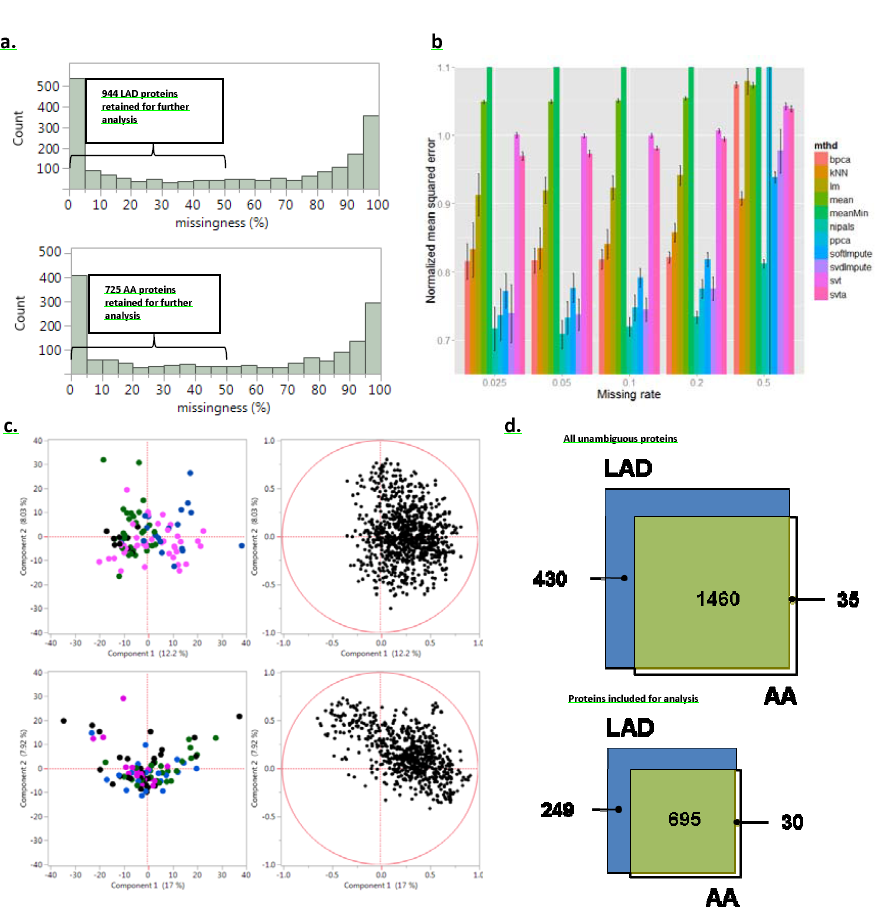
Data missingness, imputation, batch affects and anatomic overlap, **a.**) Distribution of missingness for the LAD and AA data. The most common value for missingness was 0% in both the LAD and AA. However, there was a range of missingness including some unambiguously identified protein groups that were only present in a minority of samples. Missingness encoded models did not identify any proteins with >05% missingness that were significantly associated with disease status, b.) Using proteins with complete data (0% missingness) data sets simulating various rates (2.5%-50.0%), and types (random, inversely proportional with mean signal intensity) of missingness were used to test 11 different matrix completion methods. Based on the low normalized mean squared error, the NPIALS method was selected for imputation, c.) PCA analysis with color encoding for different protein extraction batches did not identify important batch effects or extreme outliers, d.) Proportional Venn diagrams indicating the overlap in identified and analyzed proteins in the LAD and AA territories.

